# Androgen receptor-mediated assisted loading of the glucocorticoid receptor modulates transcriptional responses in prostate cancer cells

**DOI:** 10.1101/2024.11.15.623719

**Authors:** Johannes Hiltunen, Laura Helminen, Niina Aaltonen, Kaisa-Mari Launonen, Hanna Laakso, Marjo Malinen, Einari A. Niskanen, Jorma J. Palvimo, Ville Paakinaho

## Abstract

Steroid receptors are involved in a wide array of crosstalk mechanisms that regulate diverse biological processes, with significant implications in diseases, particularly in cancers. In prostate cancer, indirect crosstalk between androgen receptor (AR) and glucocorticoid receptor (GR) is well-documented, wherein AR suppression by antiandrogen therapy leads to elevated GR levels, enabling GR to compensate for and replace AR signaling. However, the existence and impact of direct chromatin crosstalk between AR and GR in prostate cancer have remained elusive. Our genome-wide investigations reveal that AR activation significantly expands GR chromatin binding. Mechanistically, AR induces remodeling of closed chromatin sites, facilitating GR binding to inaccessible sites. Importantly, coactivation of AR and GR results in distinct transcriptional responses at both the cell population and single-cell levels. Intriguingly, pathways affected by these transcriptional changes are generally associated with improved patient survival. Thus, the direct crosstalk between AR and GR yields markedly different outcomes from the known role of GR in circumventing AR blockade by antiandrogens.

## INTRODUCTION

Chromatin binding of transcription factors (TFs) to regulatory elements at enhancers plays a crucial role in shaping the development and function of cells by driving specific gene expression patterns (Field and Adelman 2020; Jindal and Farley 2021). The regulatory capabilities of TFs stem from their ability to recruit coregulators that influence both enhancer and RNA polymerase activity (Bell et al. 2024). These effects may involve alterations in chromatin accessibility or post-translational modifications of chromatin-associated proteins. Additionally, the changes initiated by one TF at an enhancer, such as increasing chromatin accessibility, can subsequently facilitate the binding of other TFs to the same enhancer (Swinstead et al. 2018; Hiltunen et al. 2024). This cooperative action among TFs, known as TF crosstalk, further broadens the cell’s gene expression patterns. Moreover, TF cooperation is recognized as a hallmark of active enhancers (Rao et al. 2021). TF crosstalk can occur through a variety of mechanisms, either directly at the same enhancer or indirectly through progressive changes in gene expression. Direct crosstalk may involve physical protein-protein interactions between TFs or can occur independently of such interactions (Rao et al. 2021; Swinstead et al. 2018; Jindal and Farley 2021).

The critical role of enhancers in regulating gene expression is underscored by their frequent association with a wide range of human diseases, including cancers, metabolic disorders, and monogenic diseases (Zaugg et al. 2022). In particular, enhancer dysfunction is commonly observed in cancers, where it can drive disease progression through both genetic and epigenetic mechanisms. While initial studies emphasized the role of DNA mutations within enhancers in tumorigenesis (Sur et al. 2012), cancers can also arise in the absence of driver mutations through epigenetic alterations at enhancers (Parreno et al. 2024). These epigenetic changes can result in the aberrant activation of oncogenic signaling pathways. Interestingly, tumorigenesis-driven enhancer activation can be linked to alterations in TF crosstalk (Rogerson et al. 2019), highlighting that cooperative TF interactions at enhancers are a key component of enhancer dysfunction. At the same time, TF crosstalk at enhancers can also help sustain the tumor-suppressive functions of specific TFs, where disruption of these interactions may contribute to increased tumorigenesis. This mechanism is particularly prevalent among steroid receptors — a subclass of the nuclear receptor superfamily — including glucocorticoid (GR), estrogen (ER), and androgen (AR) receptors, which can exert either oncogenic or tumor-suppressive effects depending on the context of their enhancer interactions (De Bosscher et al. 2020; Paakinaho and Palvimo 2021).

GR’s ubiquitous expression enables it to crosstalk with several other nuclear receptors in various cancers, particularly with ER in breast cancer (Swinstead et al. 2018; De Bosscher et al. 2020; Paakinaho and Palvimo 2021). An early study revealed that GR enhances ER’s chromatin binding by modulating chromatin accessibility, a process termed as the assisted loading mechanism (Voss et al. 2011). In this mechanism, the initiating factor (*i.e.*, GR) facilitates the binding of a secondary factor (*i.e.*, ER) to chromatin (Swinstead et al. 2018; Hiltunen et al. 2024). The key hallmarks of assisted loading include context dependency, the involvement of ATP-dependent chromatin remodeling complexes, and short-lived chromatin interactions. The latter allows both TFs to bind the same site without competition (Voss et al. 2011). Notably, in breast cancer, the GR-ER crosstalk has been found to play a protective role, with the loss or lack of this interaction contributing to tumorigenesis (Prekovic et al. 2023; Swinstead et al. 2018). However, in endometrial cancer, the ER-GR interaction is detrimental and associated with worse outcomes for patients (Vahrenkamp et al. 2018). This context-specific behavior suggests that additional cell type-specific TFs and coregulators likely influence the nature and outcome of steroid receptor crosstalk in different cancers.

Prostate cancer (PCa) progression is primarily driven by AR signaling, making antiandrogens the cornerstone of treatment for patients (Bolis et al. 2021; Paakinaho and Palvimo 2021). While AR remains the principal therapeutic target in PCa, other TFs, notably GR and FOXA1, play critical roles in disease progression and the emergence of antiandrogen resistance (Bolis et al. 2021; Hiltunen et al. 2024; Sahu et al. 2011). The ability of these TFs to interact and cooperate (Sahu et al. 2013; Swinstead et al. 2018; Helminen et al. 2024) underscores the importance of understanding their coordinated actions in influencing PCa progression and therapeutic outcomes. In contrast to the cooperative interaction between ER and GR in breast cancer, research on GR and AR crosstalk in PCa has predominantly focused on their indirect interactions (Helminen et al. 2024; Hiltunen et al. 2024; Arora et al. 2013). This indirect crosstalk stems from AR’s repression of *NR3C1* (the gene encoding GR). Antiandrogen therapy relieves this repression, leading to an increase in GR expression, allowing GR to partially replace AR on chromatin. Although this replacement is incomplete, it is sufficient for GR to regulate critical AR target genes, facilitating drug resistance and cancer progression (Arora et al. 2013; Helminen et al. 2024). Despite the significance of this indirect crosstalk, direct cooperation between AR and GR on chromatin remains elusive. Direct crosstalk is particularly significant here because AR and GR binding motifs — known as androgen (ARE) and glucocorticoid response element (GRE) — contain practically identical DNA sequence (Chen et al. 2024; Kregel et al. 2020). Consequently, AR and GR are intuitively expected to compete for chromatin binding. Moreover, direct crosstalk may yield distinct outcomes, similar to what we have recently observed in the interaction between GR and the pioneer factor FOXA1 (Huttunen et al. 2024; Helminen et al. 2024; Hiltunen et al. 2024). Like AR, FOXA1 indirectly crosstalks with GR by repressing *NR3C1* expression, but it also directly assists GR’s chromatin binding in a context-dependent manner. From a therapeutic standpoint, selectively preserving indirect while inhibiting direct crosstalk of FOXA1 with GR may offer a more targeted approach for treating PCa. Here we sought to close the knowledge gap surrounding direct crosstalk between AR and GR in PCa cells by employing a range of genome-wide techniques.

## RESULTS

### AR activation substantially expands GR’s chromatin binding in PCa cells

Analysis of PCa patient datasets indicated that *AR* and *NR3C1* are co-expressed among patients (Supplemental Fig. S1A), and patients with high expression of both *AR* and *NR3C1* have better disease-free survival than patients with high *AR* and low *NR3C1* expression (Supplemental Fig. S1B). These datasets suggest a potential direct crosstalk between AR and GR that has a different outcome from the GR-driven antiandrogen resistance. To characterize the direct crosstalk between AR and GR, we utilized VCaP cells which endogenously express transcriptionally active wild-type AR and GR (Helminen et al. 2024; Puhr et al. 2018). To evaluate how coactivation of both receptors influences their chromatin binding profile, we exposed VCaP cells to EtOH (vehicle), Dex (dexamethasone, GR agonist), DHT (dihydrotestosterone, AR agonist), or both Dex and DHT (DexDHT for simplicity). DexDHT exposure had minimal impact on AR and GR protein levels compared to single-hormone treatments (Supplemental Fig. S1C). Reporter gene assays confirmed that GR activation occurs only in response to glucocorticoids, while AR activation is specific to androgens (Supplemental Fig. S1D). Immunofluorescence analysis further demonstrated that AR translocates to the nucleus only with DHT and DexDHT treatments (Supplemental Fig. S1E), whereas GR translocates to the nucleus exclusively with Dex and DexDHT treatments (Supplemental Fig. S1F). Based on these findings, we did not conduct GR ChIP-seq with DHT-only treatment or AR ChIP-seq with Dex-only treatment. Intriguingly, ChIP-seq analyses indicated that DexDHT treatment substantially increased GR chromatin occupancy, while AR binding remained unchanged (Fig. 1A-B, Supplemental Table S1). Since AR-binding sites (ARBs) in VCaP cells were insensitive to GR activation, we focused on the GR-binding sites (GRBs). Clustering of GRBs (see Methods for details) revealed two major clusters, C1 and C2 (Fig. 1C). Example loci of C1 and C2 are presented in Fig. 1D. GR signal intensity at C1 is increased from vehicle only by DexDHT treatment, while GR is increased after both Dex and DexDHT treatments at C2 (Fig. 1E). Moreover, DexDHT further increased GR binding compared to Dex at around half of the C2 sites (Supplemental Fig. S1G). AR signal intensity does not differ between DHT and DexDHT treatments, and AR binds to C1 and C2 regardless of GR activation (Fig. 1F). In 22Rv1 cells, PCa cell line that also express AR and GR (Puhr et al. 2018; Helminen et al. 2024), ChIP-qPCR analyses indicated that AR binds to *FKBP5* and *KLK3* loci regardless of GR activation, whereas GR binding to *FKBP5* loci is enhanced by DexDHT treatment (Supplemental Fig. S2A). Moreover, ChIP-qPCR analyses in PC3 cells stably expressing AR (PC3-AR) (Kaikkonen et al. 2013) (Supplemental Fig. S2B) indicated that DexDHT treatment enhanced GR binding to *FKBP5* loci without influencing AR’s DHT-induced binding (Supplemental Fig. S2C).

**Figure 1.**
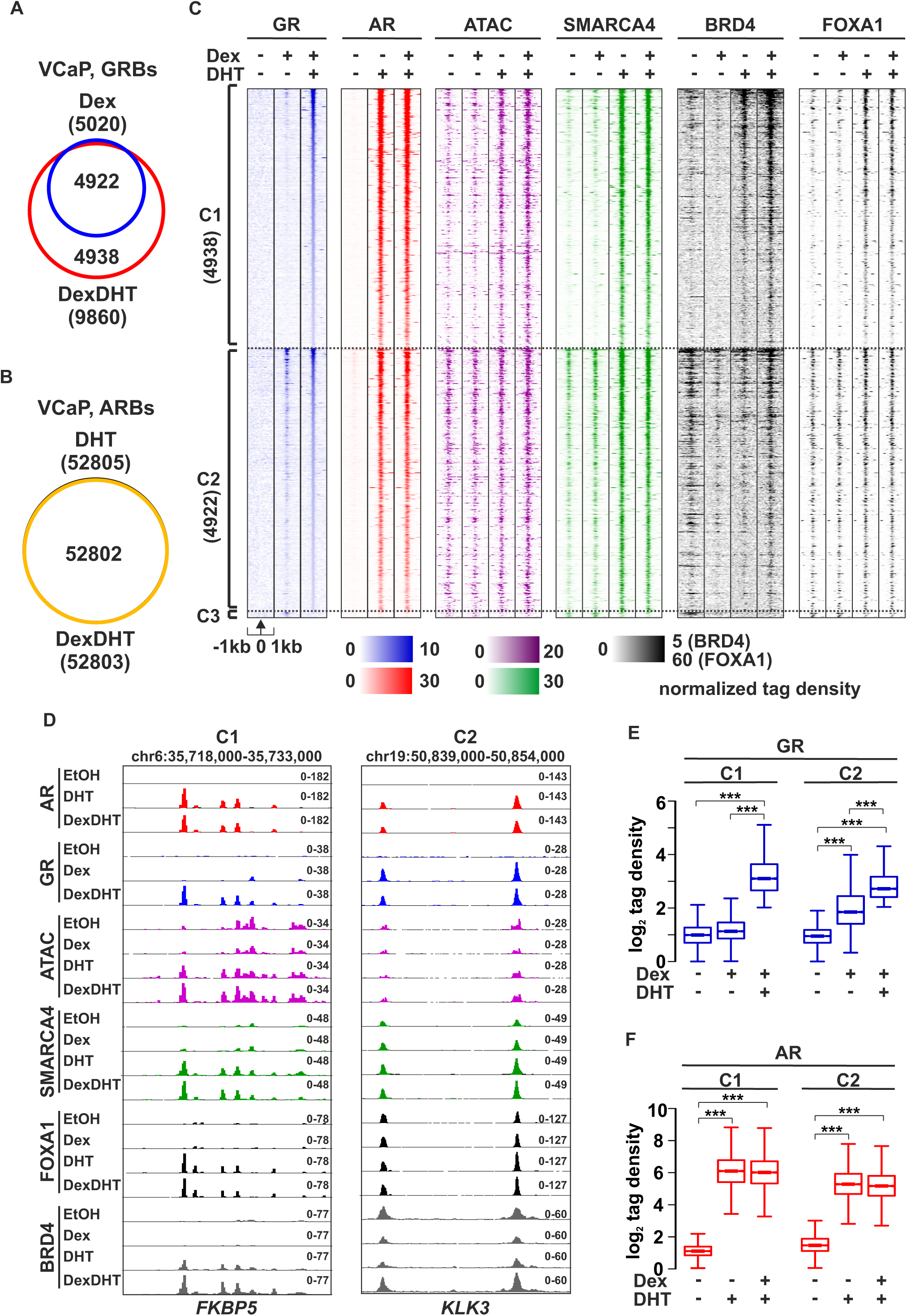
Activation of AR assists the loading of GR to chromatin. (A) Venn diagrams depict the overlap of GR binding sites (GRBs) in Dex (blue circle) and DexDHT (red circle) treated VCaP cells. (B) Venn diagrams depict the overlap of AR binding sites (ARBs) in DHT (orange circle) and DexDHT (black circle) treated VCaP cells. (C) GR ChIP-seq, AR ChIP-seq, ATAC-seq, SMARCA4 ChIP-seq, BRD4 ChIP-seq, and FOXA1 ChIP-seq profiles at C1, C2, and C3 sites in VCaP cells. C1 represents AR assisted GRBs (DexDHT/Dex: FDR<0.05, FC>2), C2 represents stable GRBs, and C3 represents AR-decreased GRBs (Dex/DexDHT: FDR<0.05, FC>2) upon the respective treatments. Each heatmap represents ± 1 kb around the center of the GR peak. Binding intensity (tags per bp per site) scale is noted below on a linear scale. (D) Genome browser track examples of AR, GR, SMARCA4, FOXA1 and BRD4 ChIP-seq, and ATAC-seq at (left) *FKBP5* and (right) *KLK3* loci in VCaP cells. (E-F) Box plots represent the normalized log_2_ tag density of (E) GR ChIP-seq and (F) AR ChIP-seq at the indicated sites. Statistical significance calculated using One-way ANOVA with Bonferroni *post hoc* test. All heatmaps, box plots, and genome browser tracks are normalized to a total of 10 million reads. ***, *p* < 0.001.

While C1 and C2 displayed no major differences in their genomic localization (Supplemental Fig. S2D), motif analyses indicated that C2 sites harbor significantly more GATA2-4, FOXA1 and HOXB13 motifs compared to C1 sites (Fig. 2A-B, Supplemental Table S2). This was concordant with the significantly higher GATA2, FOXA1 and HOXB13 motif scores at C2 compared to C1 (Supplemental Fig. S2E). In comparison, C1 sites harbored more NKX3.1, PR and half ARE motifs with significantly higher motifs scores (Fig. 2A-B, Supplemental Fig. S2E, Supplemental Table S2). While C1 and C2 harbored roughly the same percentage of ARE and GRE motifs (Fig. 2A-B), the motif scores were significantly increased in C1 compared to C2 (Supplemental Fig. S2E). Altogether, this indicates that GRBs that appear only after AR activation (C1 sites) harbor fewer pioneer factor motifs and higher number of less stringent hormone response elements (HRE, *i.e.,* PR or half ARE) (Supplemental Fig. S2F). In addition, the stringent HREs (*i.e.,* ARE or GRE) more closely resemble canonical motif form at C1 sites. These results suggest that AR has an active role at C1 sites. Finally, HRE-positive sites typically contain at least two full HRE (*i.e.,* ARE, GRE or PR) motifs (Fig. 2C), with C1 sites harboring more multi-HRE sites. This demonstrates that AR and GR can simultaneously occupy the same enhancer regions, supporting the idea that these receptors do not compete for binding at these sites. Consistent with this, co-immunoprecipitation (co-IP) analysis revealed no direct interaction between AR and GR (Supplemental Fig. S3A). However, sequential (re)ChIP assays demonstrated that AR and GR can co-occupy the *FKBP5* locus upon DexDHT treatment (Supplemental Fig. S3B-E). Notably, this type of direct crosstalk, where AR activation facilitates GR binding, has been characterized as assisted loading (Swinstead et al. 2018). Consequently, the cooperative interaction between AR and GR fundamentally differs from the known indirect crosstalk observed in antiandrogen-exposed PCa cells (Helminen et al. 2024). This is further supported by the limited (∼28%) overlap between C1 and antiandrogen-induced (ENZ-UP) GRBs, while showing substantial (∼60%) overlap between C2 and ENZ-UP GRBs (Fig. 2D).

**Figure 2.**
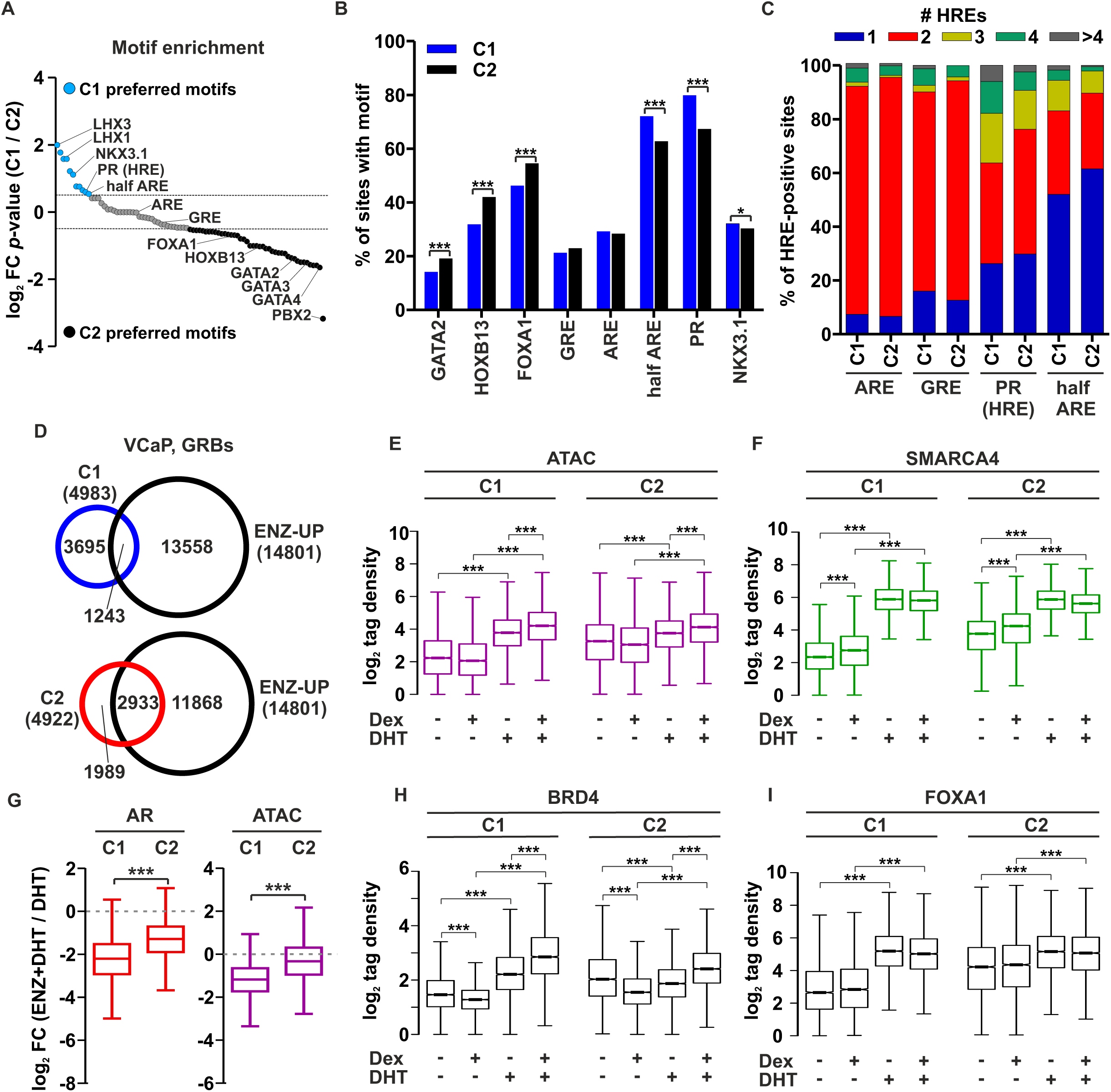
AR remodels chromatin accessibility thereby assisting the binding of GR. (A) Scatter plot represents the log_2_ fold change (FC) of motif enrichment *p*-value between C1 and C2 sites. Distinct C1 preferred (light blue) and C2 preferred (black) motifs are indicated. Grey dashed lines represent ±0.5 log_2_ FC. (B) Percentage of GATA2, HOXB13, FOXA1, GRE, ARE, half ARE, PR, and NKX3.1 motif enrichment at C1 (blue bars) and C2 (black bars) peaks. Statistical significance calculated with Chi-squared test. (C) Percentage of HRE-positive C1 and C2 sites with one (blue), two (red), three (yellow), four (green), or more than four (grey) ARE, GRE, PR, and half ARE motifs. (D) Venn diagrams depict the overlap of (upper) C1 GRBs (blue circle) and antiandrogen-induced GRBs (ENZ-UP) (black circle), and (lower) C2 GRBs (red circle) and ENZ-UP (black circle) in VCaP cells. (E-F) Box plots represent the normalized log_2_ tag density of (E) ATAC-seq and (F) SMARCA4 ChIP-seq at the indicated sites. Statistical significance calculated using One-way ANOVA with Bonferroni *post hoc* test. (G) Box plots represent the log_2_ FC of AR ChIP-seq (left) and ATAC-seq (right) in DHT treated VCaP cells in the absence or presence of antiandrogen. Grey dashed line represents the absence of change. Statistical significance calculated with unpaired *t*-test. (H-I) Box plots represent the normalized log_2_ tag density of (H) BRD4 ChIP-seq and (I) FOXA1 ChIP-seq at the indicated sites. Statistical significance calculated using One-way ANOVA with Bonferroni *post hoc* test. All box plots are normalized to a total of 10 million reads. ***, *p* < 0.001.

The ability of the initiator TF (*i.e.*, AR) that is assisting the binding of the secondary TF (*i.e.*, GR) to change chromatin accessibility can be considered as a hallmark of assisted loading (Voss et al. 2011; Swinstead et al. 2018). To evaluate if chromatin accessibility could explain the C1 and C2 differences, we conducted ATAC-seq and chromatin remodeler SMARCA4 ChIP-seq from VCaP cells exposed to EtOH, Dex, DHT, or DexDHT. In support of the assisted loading model, chromatin at C2 sites were already preaccessible without any treatment, whereas C1 sites were relatively inaccessible prior AR activation (Fig. 1C, 2E). This is in line with our previous results indicating GR’s capability to only bind to preaccessible sites in PCa cells (Helminen et al. 2024). Upon AR activation, the chromatin accessibility at C1 is increased which presumably enables GR binding to these sites in DexDHT exposed cells. Moreover, SMARCA4 ChIP-seq analyses corroborated ATAC-seq results, indicating that AR recruits the chromatin remodeling complex mediating the increased chromatin accessibility (Fig. 1C, 2F). In addition, AR ChIP-seq and ATAC-seq data from antiandrogen-treated cells (Helminen et al. 2024) revealed that DHT-induced AR binding and chromatin accessibility were more significantly reduced at C1 compared to C2 sites verifying the crucial role of AR (Fig. 2G).

In addition to steroid receptors, other TFs and coregulators have been indicated to contribute to assisted loading (Swinstead et al. 2016; Goldstein et al. 2017). Because of this, we also profiled the chromatin binding of one TF, FOXA1, and one coregulator, BRD4, to these sites. We chose FOXA1 and BRD4, since AR is the initiator TF, and both proteins can modulate AR signaling (Sahu et al. 2011; Asangani et al. 2014). Intriguingly, BRD4 and FOXA1 occupy C2 sites in the absence of any treatments, while their binding to C1 sites occur only after AR activation (Fig. 1C, 2H-I). While SMARCA4 and FOXA1 ChIP-seq signals did not differ between DHT and DexDHT treatments (Fig. 2F, 2I), the DHT-induced ATAC-seq and BRD4 ChIP-seq signals significantly increased when combined with Dex treatment (Fig. 2E, 2H). To analyze these results more broadly, we sorted all ARBs based on chromatin accessibility to quartiles (Q1-Q4) with Q1 being the most and Q4 the least preaccessible (Supplemental Fig. S4A, Supplemental Table S1). Intriguingly, motif analyses indicated that the more preaccessible the ARB is, the more FOXA1 motifs it contains, while the less preaccessible the ARB is the more HRE motifs it contains (Supplemental Fig. S4B, Supplemental Table S2). The analysis of all ARBs (Q1-Q4) was concordant with GRBs (C1-C2), wherein GR can bind to less accessible sites only when AR is activated, and at these sites AR increases chromatin accessibility and the recruitment of SMARCA4, FOXA1, and BRD4 (Supplemental Fig. S3C-H). In addition, at all ARBs, ATAC-seq and BRD4 ChIP-seq signals significantly increased between DHT and DexDHT treatments. Our results indicate that not only is AR assisting the binding of GR to chromatin, but it also induces the recruitment of FOXA1 and BRD4 to the sites in PCa cells. For FOXA1, reChIP assays confirmed the co-occupancy of GR and FOXA1 at the *FKBP5* locus following DexDHT treatment and at the *KLK3* locus upon Dex and DexDHT treatments (Supplemental Fig. S3B-E). Thus, GR seems not to be a mere bystander at the sites since it can increase the DHT-induced chromatin accessibility and BRD4 chromatin occupancy.

### Coactivation of AR and GR leads to synergistic expression of a subset of target genes

Because GR at AR-assisted GRBs has added effect on chromatin accessibility, we next measured if coactivation of both receptors change transcriptional regulation. For this, we performed global run-on sequencing (GRO-seq) from VCaP cells exposed to EtOH, Dex, DHT, or DexDHT to capture primary transcriptional regulatory events. Since GRO-seq measures ongoing transcription (Toropainen et al. 2016), we first evaluated the enhancer RNA (eRNA) production at intergenic C1 and C2 sites. These results were concordant with ATAC-seq results indicating that C1 sites produce very little eRNA without AR activation, whereas C2 sites were already transcriptional active (Fig. 3A, Supplemental Fig. S5A). AR activation substantially increased the production of eRNA at C1 sites after DHT and DexDHT treatment. Intriguingly, while GR on its own had no effect on chromatin accessibility (Fig. 2E), it increased eRNA production at C2 sites upon Dex treatment (Fig. 3A, Supplemental Fig. S5A), indicating that the GR can activate these enhancers. Furthermore, DHT-induced chromatin accessibility did not consistently correlate with increased eRNA production at C1 sites, and *vice versa* (Supplemental Fig. S5B). This suggests that ATAC-seq and GRO-seq capture both overlapping and distinct regulatory effects. Subsequently, using GRO-seq data we defined differentially hormone-regulated genes (see Methods for details). While there are only tens of Dex-regulated genes, DHT exposure led to differential expression of well over thousand genes (Fig. 3B, Supplemental Table S3). This is comparable number of androgen-regulated genes in VCaP cells defined in previous GRO-seq experiments (Toropainen et al. 2016). Intriguingly, coactivation of both AR and GR led to a substantial increase of hormone-regulated genes compared to DHT only treatment (Fig. 3B), and only a small proportion of genes lost their hormone-regulation upon DexDHT treatment. In support, RT-qPCR analyses performed in PC3-AR cells indicated that the expression of *FKBP5* and *RASD1* was induced by DexDHT treatment compared to DHT treatment, while *TMPRSS2* and *SGK1* showed no such induction (Supplemental Fig. S5C).

**Figure 3.**
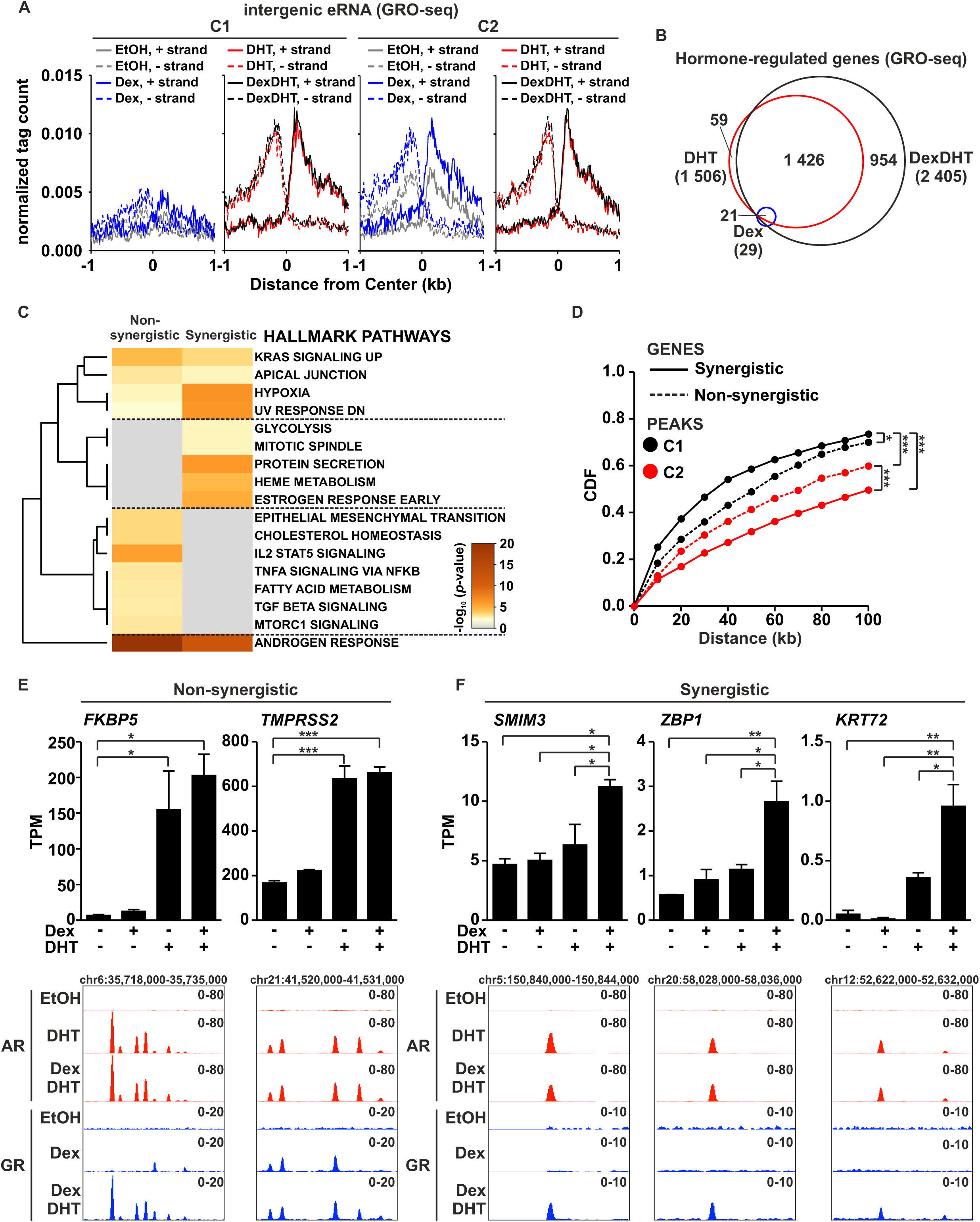
Coactivation of AR and GR synergistically induces a subset of target genes. (A) Aggregate plots represent the eRNA intensity of GRO-seq at intergenic C1 and C2 peaks in EtOH (grey line), Dex (blue line), DHT (red line), and DexDHT (black line) treated VCaP cells. GRO-seq tags represented in strand-specific manner with separated plus (solid line) and minus (dashed line) strand. Each aggregate plot represents ± 1 kb around the center of the binding sites. (B) Venn diagrams depict the overlap of Dex (blue circle), DHT (red circle), and DexDHT (black circle) regulated genes in VCaP cells. (C) Hallmark Gene Sets pathway analysis of synergistic and non-synergistic genes in VCaP GRO-seq data. Color scale represents - log_10_ *p*-value. (D) Association of non-synergistic (dashed line) and synergistic (solid line) genes with C1 (black color) and C2 (red color) peaks. Data is represented as cumulative distribution function (CDF). Statistical significance calculated with One-way ANOVA with Bonferroni *post hoc* test. (E) Example of non-synergistic *FKBP5* and *TMPRSS2* genes. (F) Example of synergistic *SMIM3*, *ZBP1* and *KRT72* genes. Bar graphs (upper) depict target gene expression using GRO-seq tags (TPM values). Genome browser tracks (lower) depict chromatin binding of AR and GR. Statistical significance calculated with One-way ANOVA with Bonferroni post hoc test. All genome browser tracks are normalized to a total of 10 million reads. *, *p* < 0.05; **, *p* < 0.01; ***, *p* < 0.001.

To define the difference between DHT and DexDHT treatments more closely, we focused on the induced genes in DHT and DexDHT exposed VCaP cells. We defined the genes either as DHT- and DexDHT-induced (non-synergistic) or synergistically DexDHT-induced (synergistic) genes (see Methods for details) (Supplemental Fig. S5D, Supplemental Table S4). For the former, the induction of hormone-regulated genes is similar between DHT and DexDHT treatment. For the latter, coactivation of AR and GR leads to synergistic induction of hormone-regulated genes compared to single hormone treatments. Definition of synergistically induced genes was implemented as described by Goldberg et al. (Goldberg et al. 2022). Of the hormone-induced genes, 47% were synergistically regulated (Supplemental Fig. S5E). Intriguingly, pathway analyses indicated clear differences between non-synergistic and synergistic genes (Fig. 3C). While androgen response was regulated by both sets of genes, synergistically induced genes regulated several distinct pathways, such as glycolysis, mitotic spindle, and estrogen response early. The non-synergistically induced genes were enriched at pathways commonly associated with androgen-regulated genes, such as cholesterol homeostasis, fatty acid metabolism, and MTORC1 signaling (Lafront et al. 2024; Helminen et al. 2024). Moreover, C1 GRBs, where AR assists the binding of GR, were significantly more closely associated with synergistic than non-synergistic genes (Fig. 3D). The opposite was observed with C2 GRBs, where GR binds primarily regardless of AR activation. As representative examples, the *FKBP5* and *TMPRSS2* expression do not show synergistic induction, and their regulatory regions contain C2 GRBs (Fig. 3E). It is important to note that both loci contain also C1 GRBs. In comparison, *SMIM3*, *ZBP1*, and *KRT72* are examples of synergistically induced genes, and their regulator regions contain at least one prominent C1 GRB (Fig. 3F). Our results indicate that coactivation of GR and AR in PCa cells leads to synergistic regulation of distinct gene pathways whose regulation is linked to AR-assisted GRBs.

### Distinct set of PCa cells respond to AR and GR coactivation at single-cell transcriptome level

Since coactivation of AR and GR lead to synergistic transcriptional regulation at cell population level, we aimed to address this also at single cell level. For this, we performed single-cell RNA-seq (scRNA-seq) from VCaP cells exposed to EtOH, Dex, DHT, or DexDHT. After filtering (see Methods for details), uniform manifold approximation and projection (UMAP) clustering of all samples indicated that one cluster of cells was formed primarily by EtOH and Dex treated cells, while DHT and DexDHT treated cells formed another cluster (Fig. 4A). This suggests that Dex on its own has only a limited transcriptional response at single VCaP cell level. Integration of single cells from all four treatment groups resulted in 16 subpopulations of cells (SP0-SP15) (Fig. 4B). Single cells derived from EtOH and Dex treated samples were in SP2, SP3, SP5, SP6, SP7, SP9, and SP12, while single cells derived from DHT- and DexDHT-treated samples were in SP0, SP1, SP4, SP8, SP10, SP13, and SP14 (Supplemental Fig. S6A). The mixture of single cells from all treatments were in SP11 and SP12. As examples, *FKBP5* and *TMPRSS2* have high mRNA expression and percent expressed cells in the indicated DHT and DexDHT subpopulations (Fig. 4C-D).

**Figure 4.**
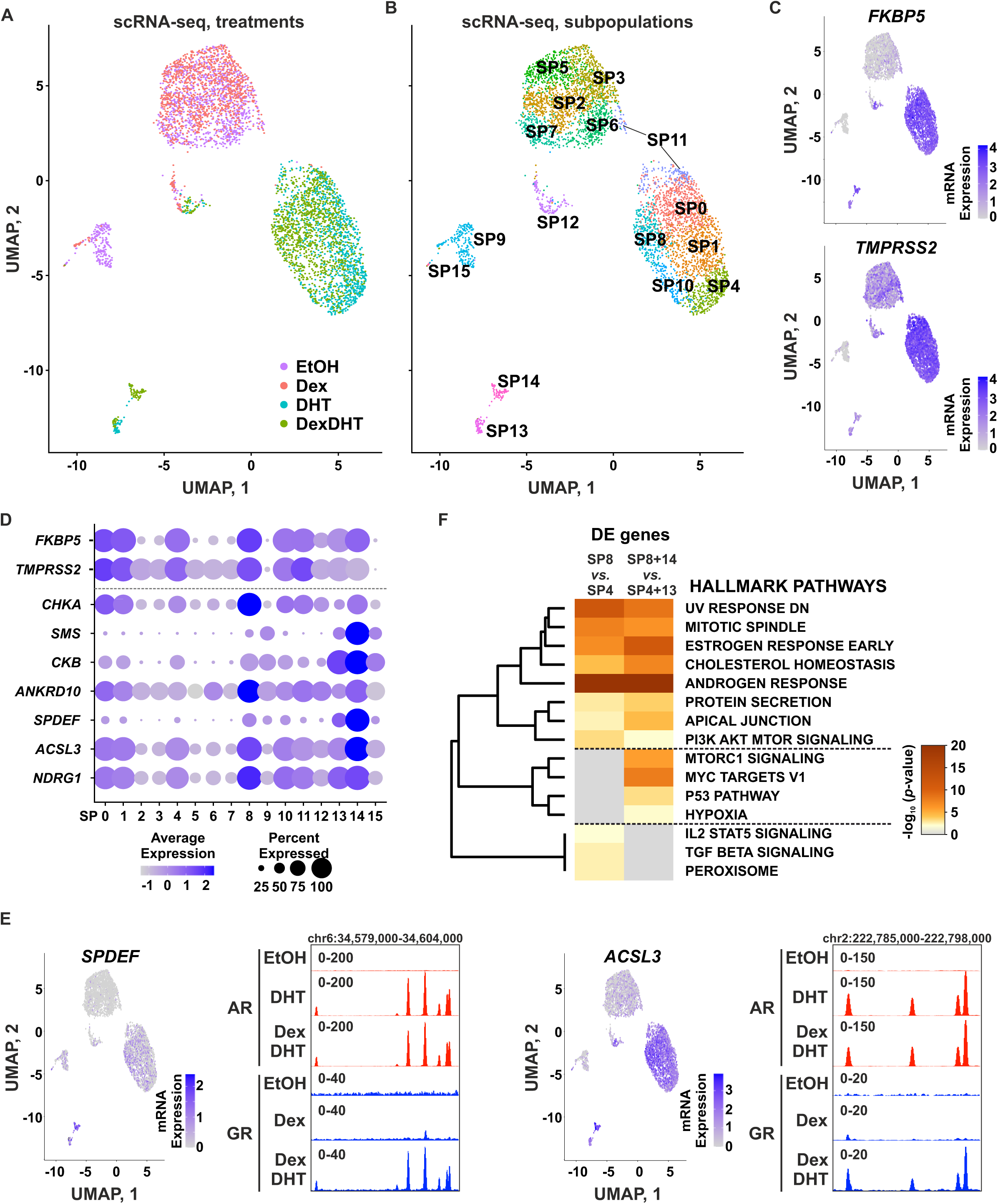
Coactivation of AR and GR influences a distinct population of single cells. (A) Uniform Manifold Approximation and Projection (UMAP) of scRNA-seq from EtOH (purple dots), Dex (red dots), DHT (blue dots), and DexDHT (green dots) treated VCaP cells. (B) UMAP clustering from the scRNA-seq with subpopulations 0-15 (SP0-SP15) indicated with distinct color and label. (C) UMAP of (upper) *FKBP5* and (lower) *TMPRSS2* expression at the single cell subpopulations. Intensity of the color depicts the mRNA expression (log1p normalized). (D) Dot plot of the expression of selected genes in the single cell subpopulations. The size of the circle depicts the percentage of cells that express the gene. Intensity of the color depicts the average expression. (E) UMAP of gene expression at the single cell subpopulations and genome browser tracks of chromatin binding of AR and GR for (left) *SPDEF* and (right) *ACSL3*. (F) Hallmark Gene Sets pathway analysis of differentially expressed (DE) genes between SP8 and SP4, and between SP8+14 and SP4+13 in scRNA-seq data. Color scale represents −log_10_ *p*-value. All genome browser tracks are normalized to a total of 10 million reads.

Examination of the DHT and DexDHT treatments indicated that single cells in certain subpopulations have differential responses to the treatments. SP4 and SP13 primarily consist of single cells from DHT treatment, while SP8 and SP14 primarily consist of single cells from DexDHT treatment (Fig. 4A-B, Supplemental Fig. S6A). As examples, *SPDEF*, *ACSL3*, and *NDRG1* (as well as other targets) have higher average expression and percent expressed cells in SP8 and SP14 compared to SP4 and SP13 (Fig. 4D). Moreover, when examining treatments rather than subpopulations, these genes are the most highly expressed in DexDHT treated cells (Supplemental Fig. S6B). These example genes also harbor prominent C1 GRBs at their regulatory regions, indicating that AR-assisted loading of GR modulates their gene expression (Fig. 4E, Supplemental Fig. S6C). Intriguingly, genes differentially regulated between single cell subpopulations of DHT and DexDHT treatments (SP8 *vs.* SP4, or SP8+14 *vs.* SP4+13), were enriched at pathways such as estrogen response early, cholesterol homeostasis, and apical junction (Fig. 4F). Moreover, 40% of the differentially regulated genes between DHT and DexDHT in scRNA-seq were associated with C1 GRBs (Supplemental Table S4). The genes with C1 GRB were associated with hypoxia and IL2 STAT5 signaling pathways while the genes without C1 GRB were associated with pathways such as cholesterol homeostasis and MTORC1 signaling (Supplemental Fig. S6D). Androgen response and estrogen response early pathways were enriched with genes with or without C1 GRB. In concordance with cell population level analyses, our single cell level results indicated that coactivation of GR and AR leads to regulation of distinct pathways and their regulation can be linked to AR-assisted GRBs.

### Estrogen-related receptor gamma and GR are cooperatively associated with beneficial PCa patient outcomes

To assess the biological impact of AR and GR coactivation, we measured proliferation rates in VCaP cells. While Dex treatment alone had no effect, DHT treatment significantly stimulated VCaP cell proliferation (Fig. 5A). Interestingly, the combined DexDHT treatment substantially attenuated the DHT-induced proliferation, suggesting that the distinct pathways activated by DexDHT (Fig. 3C, 4F, Supplemental Fig. S6D) may restrict DHT-driven PCa cell growth. To evaluate potential downstream TFs mediating these effects, we analyzed synergistic genes identified by GRO-seq and SP8+14 genes from scRNA-seq. Notably, ER (*ESR1*, ERα) emerged as a candidate regulator of DexDHT-induced effects (Supplemental Fig. S7A). ER has recently been implicated with PCa disease progression with known roles in regulating metabolic pathways (Lafront et al. 2024). Correspondingly, both estrogen response early and metabolic pathways are associated with DexDHT effects (Fig. 3C, 4F, Supplemental Fig. S6D). Additionally, in breast cancer, both AR and GR can influence ER signaling, resulting in survival benefits (Paakinaho and Palvimo 2021), suggesting that GR potentially modulates AR-regulated estrogen signaling in PCa. However, comparison of the estradiol-regulated transcriptome (Lafront et al. 2024) with differential DexDHT genes indicated minimal overlap (Supplemental Fig. S7B). While C1 GRBs are present at the *ESR1* locus (Supplemental Fig. S7C), *ESR1* shows minimal expression and no response to hormone treatments across GRO-seq, scRNA-seq, and RT-qPCR analyses (Supplemental Fig. S7D-I). These results suggest that ER is unlikely to play a significant role in the synergistic effects observed with AR and GR coactivation.

**Figure 5.**
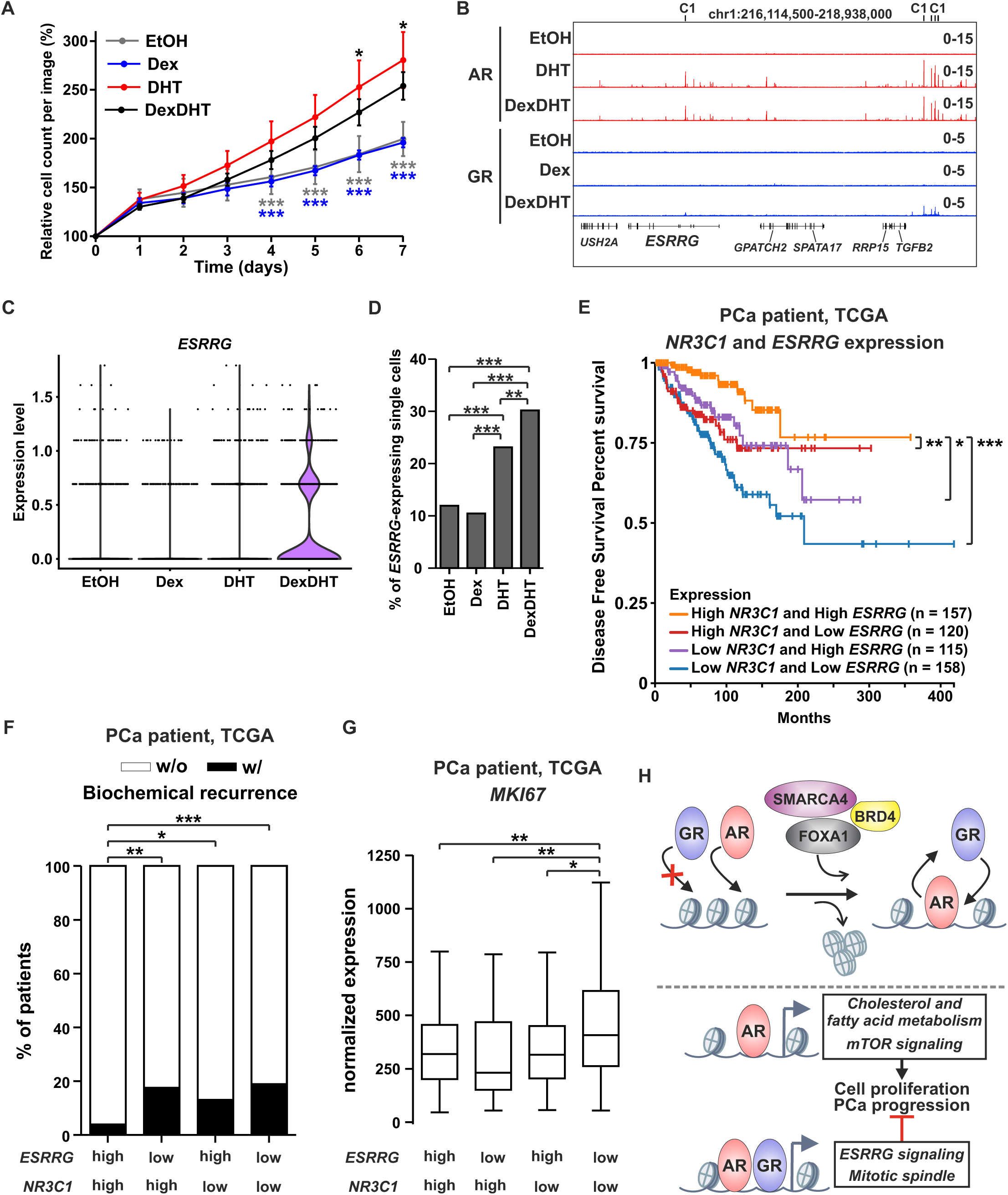
High expression of *NR3C1* and *ESRRG* are associated with the best patient outcomes. (A) Relative cell proliferation from EtOH (grey line), Dex (blue line), DHT (red line), and DexDHT (black line) treated VCaP cells. Data is normalized to day 0 and represents the mean ±SD of 8-10 images per day. Statistical significance calculated with Two-way ANOVA with Bonferroni *post hoc* test, and comparisons between DHT with other treatment are shown denoted with colored stars. (B) Genome browser track depicts chromatin binding of AR and GR at the *ESRRG* locus. (C) Violin plot represents the single cell expression level of *ESRRG* in EtOH, Dex, DHT, and DexDHT treated VCaP cells. (D) Percentage of *ESRRG*-expressing single cells in EtOH, Dex, DHT, and DexDHT treated VCaP cells. Statistical significance calculated with Chi-squared test. (E) Disease-free survival of TCGA PCa patients with combination of high and/or low expression levels of *NR3C1* and *ESRRG*. Statistical significance calculated with log-rank test. (F) Percentage of TCGA PCa patients with or without biochemical recurrence event divided based on high *NR3C1* and high *ESRRG* (*n*=124), high *NR3C1* and low *ESRRG* (*n*=123), low *NR3C1* and high *ESRRG* (*n*=123), or low *NR3C1* and low *ESRRG* (*n*=123) expression. White bars represent no biochemical recurrence event (w/o), and black bars represent biochemical recurrence event (w/). Statistical significance calculated with Chi-squared test. (G) Box plots represent normalized proliferation marker *MKI67* expression of TCGA PCa patients with high *NR3C1* and high *ESRRG* (*n*=124), high *NR3C1* and low *ESRRG* (*n*=123), low *NR3C1* and high *ESRRG* (*n*=123), or low *NR3C1* and low *ESRRG* (*n*=123) expression. (H) Model of AR assisting the loading of GR to chromatin through modulation of chromatin accessibility and enhancer activation (upper), and expansion of AR-regulated transcriptome by GR that leads to restriction of cell proliferation and PCa progression (lower). Statistical significance calculated using One-way ANOVA with Bonferroni *post hoc* test. All genome browser tracks are normalized to a total of 10 million reads. *, *p* < 0.05; **, *p* < 0.01; ***, *p* < 0.001.

In addition to ER, estrogen-related receptor gamma (*ESRRG*, ERRγ) appears to play a key role in regulating DexDHT-associated metabolic and estrogen response pathways in PCa cells (Audet-Walsh et al. 2017). Analysis of GRO-seq and scRNA-seq data revealed that *ESRRG* is expressed at higher levels than *ESR1* (Supplemental Fig. S7D, S7G). Notably, the *ESRRG* locus contains both intronic and intergenic C1 GRBs (Fig. 5B), and its expression is elevated in DexDHT-treated single cells as well as in DexDHT-responsive single-cell subpopulations (SP8+14) (Fig. 5C, Supplemental Fig. S7E-G). Furthermore, the proportion of single cells expressing *ESRRG* is highest in DexDHT-treated samples (Fig. 5D). Additional RT-qPCR analyses showed that both DHT and DexDHT treatments increase *ESRRG* expression in VCaP and PC3-AR cells (Supplemental Fig. S7I). Together, these findings suggest that AR-mediated assisted loading of GR enhances *ESRRG* expression in a subset of cells, potentially contributing to the observed changes in cell growth.

To further investigate the relationship between *ESRRG* and steroid receptors in PCa, we analyzed data from TCGA patient datasets. High *ESRRG* expression was associated with improved disease-free survival in PCa patients, a benefit not observed with high *ESR1* expression (Supplemental Fig. S7J). Intriguingly, stratifying patients based on combined expression of *NR3C1* and *ESRRG* revealed that high levels of both were linked to the most favorable survival outcomes (Fig. 5E). This effect was not seen with the expression of *AR* and *ESRRG* (Supplemental Fig. S7K). However, the survival of PCa patients with high *AR* expression was significantly improved with high expression of *ESRRG*. The positive association between GR and ERRγ extends beyond survival outcomes. PCa patients with high expression of both *NR3C1* and *ESRRG* showed significantly lower rates of biochemical recurrence compared to those with low levels of either factor (Fig. 5F). Moreover, expression of the mitosis marker *MKI67* (encoding for Ki-67) (Uxa et al. 2021) was markedly higher only in patients with low expression of both *NR3C1* and *ESRRG* (Fig. 5G). These findings suggest that AR-assisted loading of GR results in the synergistic regulation of specific gene sets, including *ESRRG*-mediated signaling. Furthermore, these data imply that GR modulation of AR’s transcriptional activity, combined with co-expression of GR and ERRγ, is associated with better clinical outcomes for PCa patients.

## DISCUSSION

The complex interplay between AR and GR in promoting antiandrogen resistance through indirect mechanisms has been well-explored by us and others (Desai et al. 2024; Arora et al. 2013; Helminen et al. 2024; Puhr et al. 2018). However, relatively little is known about the direct cooperation between AR and GR in PCa cells. In contrast, direct crosstalk between GR and ER, as well as ER and AR, has been well-characterized in breast cancer (Hickey et al. 2021; Paakinaho and Palvimo 2021; Swinstead et al. 2018; Prekovic et al. 2023). Notably, direct crosstalk is not limited to steroid receptors alone — other TFs like FOXA1, NFκB, and CREB have also been shown to cooperate directly with AR or GR in PCa cells and beyond (Sahu et al. 2011; Swinstead et al. 2016; Rao et al. 2011; Malinen et al. 2017; Goldberg et al. 2022). Over a decade ago, a study suggested potential direct AR-GR crosstalk in PCa cells by showing that coactivation of both receptors elicited a distinct transcriptional response compared to activation of either receptor alone (Sahu et al. 2013). However, these studies were conducted in engineered PCa cells. Given that crosstalk mechanisms can differ between engineered and native cell environments — such as that between GR and FOXA1 (Sahu et al. 2011; Helminen et al. 2024)— it is crucial to study these effects in a more native context.

Our genome-wide analyses in native VCaP cells revealed that AR facilitates GR binding to previously inaccessible chromatin sites by recruiting coregulators that increase chromatin accessibility and activate enhancers (Fig. 5H). This process, known as assisted loading, occurs when an initiator TF enables the chromatin binding of a secondary TF (Swinstead et al. 2018; Voss et al. 2011). Interestingly, AR not only facilitates GR binding but also promotes the binding of FOXA1 to the same sites, indicating that assisted loading may extend to multiple secondary TFs simultaneously, rather than being limited to one per initiator TF. This suggests that AR may create TF “hotspots”, similar to those driving cell differentiation (Siersbæk et al. 2014), potentially establishing key regulatory hubs in PCa. This is supported by the fact that optimal recruitment of coregulators and TFs — especially within phase-separated condensates — is crucial for AR-mediated enhancer activation (Chen et al. 2023; Zhang et al. 2023). Additionally, other TFs may contribute to the AR-mediated assisted loading. One potential candidate is the prostate luminal TF NKX3.1, as its motif showed a slight but significant association with AR-induced GRBs. NKX3.1 is known to interact with AR and GR in prostate cells (Launonen et al. 2021; Dutta et al. 2016), and its expression is often lost during the progression of PCa (Bolis et al. 2021; Dutta et al. 2016).

Despite AR and GR binding to the same chromatin sites upon coactivation, they exhibit no competition for binding. This lack of competition aligns with previous studies examining GR-associated assisted loading (Miranda et al. 2013; Swinstead et al. 2016; Goldberg et al. 2022). However, these studies mainly focused on TFs that bind to different motif sequences. Our findings extend this understanding by demonstrating that even the TFs sharing the same motif sequence, such as GR and AR, do not compete for binding across the genome. This lack of competition is likely due to relatively short TF residence times and the low occupancy of binding sites at any given moment (Wagh et al. 2023; Voss et al. 2011). Moreover, AR-mediated recruitment of SMARCA4 to chromatin further supports this effect. SWI/SNF chromatin remodeler, with SMARCA4 as the ATPase subunit, actively displaces TF from the chromatin (Li et al. 2015). This suggests that AR-induced increases in chromatin accessibility may, paradoxically, facilitate its own chromatin displacement, allowing GR to bind at the same enhancer.

Another possible explanation for the lack of competition between GR and AR is heterodimerization, a mechanism proposed more than two decades ago (Chen et al. 1997). However, GR-AR heterodimers were initially thought to exert an inhibitory effect on transcriptional activity, in contrast to the synergistic effects observed in our coactivation experiments. Moreover, we did not detect direct interaction between AR and GR. Given that GRE- and ARE-positive sites frequently contain two or more HRE sequences, it is more likely that GR and AR bind simultaneously at these sites, each occupying distinct motifs rather than forming heterodimers. This is supported by our reChIP analyses. Interestingly, canonical AREs have been linked to both oncogenic (Kregel et al. 2020) and tumor-suppressive activities (Chen et al. 2024). While ARE enrichment showed no major differences across various GRBs, AR-induced GRBs exhibited fewer pioneer factor motifs and a higher prevalence of less stringent HREs. This pattern aligns with the tumor-suppressive behavior of AREs (Chen et al. 2024), suggesting a potential role for GR in this regulatory process.

Although GR can modulate ER binding in breast and endometrial cancers (Miranda et al. 2013; Vahrenkamp et al. 2018; Prekovic et al. 2023), it remains unclear why GR does not similarly influence AR binding in PCa cells. One possible explanation is protein abundance, as AR is highly expressed in VCaP cells, whereas GR levels are relatively low (Puhr et al. 2018; Helminen et al. 2024). However, in 22Rv1 and PC3-AR cells, which express more GR and less AR than VCaP cells, GR still fails to impact AR occupancy at selected binding sites. A more likely reason for this discrepancy is the distinct chromatin landscape of PCa cells, including VCaP and 22Rv1, where GR binding is limited to preaccessible sites—a pattern not observed in other cell types (Helminen et al. 2024). This suggests that cell type-specific TFs or coregulators may either enable or restrict GR’s capacity to bind inaccessible chromatin and assist other TFs, such as AR, in their binding. Intriguingly, in U2OS osteosarcoma cells, GR binds more low-accessibility regions and recruits mediator complex subunits to chromatin more efficiently than AR (Kulik et al. 2021). This suggests that in the same cellular environment, differential coregulator recruitment may influence each receptor’s ability to access closed chromatin sites. Future chromatin proteomic studies on GR- and AR-associated partners will clarify whether this preference also occurs in PCa cells.

Although activated GR does not impact AR’s chromatin binding, it does modulate DHT-mediated transcriptional responses. At assisted sites, GR enhances DHT-induced chromatin accessibility and BRD4 occupancy, though it does not affect SMARCA4 recruitment. Notably, SMARCA4 is known to precede and facilitate the recruitment of additional coregulators to enhancers (Saha et al. 2024). Since AR increases chromatin accessibility *via* SMARCA4, enabling GR binding, it appears that GR further modulates enhancer activity through other coregulators, such as BRD4. More crucially, AR and GR coactivation results in the synergistic expression of target genes enriched in specific pathways, including mitotic spindle, protein secretion, and estrogen response early pathways. These pathways are also differentially enriched in our single-cell analyses. By contrast, non-synergistic genes are associated with pathways typically regulated by AR, such as MTORC1 signaling and cholesterol homeostasis (Lafront et al. 2024; Helminen et al. 2024). These findings suggest that the synergistic gene expression reflects a GR-mediated expansion of the AR-regulated transcriptome in PCa cells.

Our gene expression analyses suggest a prominent role for ERRγ in conjunction with GR in PCa. High co-expression of *NR3C1* and *ESRRG* is associated with improved patient survival and a reduced incidence of biochemical recurrence, aligning with findings from other studies that report protective effects of *ESRRG* expression at both the population and single-cell levels (Fan et al. 2023; Audet-Walsh et al. 2017). However, our data specifically indicated that the survival benefit conferred by ERRγ is closely linked to GR expression. The beneficial effects of ERRγ likely stem from its capacity to modulate metabolic pathways, including oxidative phosphorylation (Audet-Walsh et al. 2017). Since oxidative phosphorylation occurs in mitochondria, it is notable that GR and its splice variants are involved in regulating mitochondrial function (Morgan et al. 2016). In lung cancer cells, for example, GR activation has been shown to reduce mitochondrial respiration and downregulate cell cycle-related genes (Prekovic et al. 2021). In our PCa patient data analysis, we found that expression of the mitosis marker *MKI67* is significantly lower when either *NR3C1* or *ESRRG* is highly expressed, while low expression of both *NR3C1* and *ESRRG* is associated with elevated *MKI67* levels. This inverse relationship between GR activity and *MKI67* expression has also been observed in breast cancer (Prekovic et al. 2023). These findings suggest that the synergistic gene regulation mediated by steroid receptor-assisted loading, possibly involving ERRγ, acts to limit oxidative metabolism and suppress mitosis in PCa cells, contributing to more favorable patient outcomes.

Our ChIP-seq data revealed that AR mediates the binding of GR to multiple enhancers at the *ESRRG* locus. According to our scRNA-seq data, GR binding at these assisted enhancers leads to increased *ESRRG* expression and a higher proportion of *ESRRG*-positive single cells following DexDHT treatment. Interestingly, the induction of *ESRRG* upon AR activation in the presence or absence of activated GR is not evident in population-based assays. This suggests that AR-induced GR binding facilitates *ESRRG* expression specifically in a subset of cells. While previous studies have reported that AR represses *ESRRG* expression (Audet-Walsh et al. 2017), this discrepancy may stem from differences in the experimental models used. In our study, we employed GR-expressing VCaP and PC3-AR PCa cells, whereas the earlier study used LNCaP cells, which lack GR expression. Therefore, the activation of AR in the presence of GR could be a key factor in regulating *ESRRG* expression. Supporting this, earlier research in skeletal muscle cells showed that ERRγ can induce *NR3C1* expression (Wang et al. 2010), further highlighting the cooperative relationship between ERRγ and GR. Although our findings underscore the critical role of GR and ERRγ in PCa patient outcomes, we did not fully explore how *ESRRG* contributes to the DexDHT-mediated synergistic gene regulation. Since chromatin proteomics in VCaP cells (Launonen et al. 2021) has not identified ERRγ as a chromatin-associated partner of AR, it is likely that ERRγ exerts its effects outside the AR-GR co-occupied enhancers. To further validate the role of ERRγ, additional ChIP and gene expression analyses in PCa cells are needed.

Our exploration of the direct crosstalk between AR and GR in PCa cells suggests that GR exerts tumor-suppressive effects by modulating the AR-regulated transcriptome (Fig. 5H). This manifests as a restriction in the AR-induced cell proliferation. This contrasts sharply with GR’s well-documented oncogenic role in bypassing AR blockade by antiandrogens (Hiltunen et al. 2024). Therefore, direct AR-GR crosstalk appears to produce significantly different outcomes compared to their indirect interaction. A similar pattern can be seen in GR’s interaction with FOXA1, where FOXA1 represses GR expression through indirect crosstalk while simultaneously facilitating GR chromatin binding through direct cooperation (Huttunen et al. 2024). However, unlike AR, which assists GR loading at previously inaccessible chromatin sites, FOXA1 promotes GR occupancy at other already accessible sites. This dichotomy in GR’s function depending on the presence or absence of AR is reminiscent of its behavior in breast cancer, where GR has a protective effect when co-expressed with ER, improving patient survival. However, in ER-negative cases, GR’s expression becomes detrimental (Paakinaho and Palvimo 2021). These observations suggest that in steroid receptor-driven cancers, GR’s tumor-suppressive effects rely on the presence of AR or ER, but when these primary oncogenic steroid receptors are lost—such as in ER-negative breast cancer or antiandrogen-treated PCa — GR converts into a driver or contributor of the disease. As a result, understanding steroid receptor expression patterns could provide critical insights into the potential benefits or drawbacks of glucocorticoid treatments in cancer patients. Further research is also needed to elucidate what additional tumor-suppressive or oncogenic roles GR may play in the prostate (Hiltunen et al. 2024).

## METHODS

### Cell culture and treatments

VCaP (ATCC, #CRL-2876, RRID:CVCL_2235) and COS-1 (ATCC, #CRL-1650, RRID: CVCL_0223) cells were cultured in DMEM (Gibco, #41965039) supplemented with 10% FBS (Cytiva, #11531831) and 1 U/µl penicillin and 1 µg/ml streptomycin (Gibco, #15140122). 22Rv1 (ATCC, #CRL-2505, RRID: CVCL_1045) cells were maintained in RPMI-1640 medium (Gibco, #11875093) supplemented with 10% FBS (Gibco, #11573397), 1 U/μl penicillin, 1 μg/ml streptomycin and 2 mM L-glutamine (Gibco, #A2916801). PC3-AR (Kaikkonen et al. 2013) cells were maintained in Ham’s F12-K medium (Gibco, #21127022) supplemented with 10% FBS (Gibco, #11573397), 1 U/μl penicillin and 1 μg/ml streptomycin (Gibco, #15140122). Cells were maintained at 37°C in 5% CO_2_ conditions, and routinely checked for mycoplasma contamination. The identity of the cell lines was verified by Institute for Molecular Medicine Finland (Helsinki, Finland). For the experiments, growth medium was changed to medium supplemented with 5% charcoal-stripped FBS (Gibco, #10270106) for at least one day before the start of treatments. Cells were treated individually or simultaneously with dexamethasone (Dex) (Sigma-Aldrich, #D4902) and 5α-dihydrotestosterone (DHT) (Steraloids, #A2570-000).

### Antibodies

Primary antibodies: anti-GR (Santa Cruz Biotechnology, #sc-1003, RRID:AB_631572), anti-GR (Cell Signaling Technology, #12041, RRID:AB_2631286), anti-AR (Karvonen et al. 1997), anti-FOXA1 (Abcam, #ab23738, RRID:AB_2104842), anti-SMARCA4 (Abcam, #ab110641, RRID:AB_10861578), anti-BRD4 (Bethyl Laboratories, #A301-985A, RRID:AB_1576498), anti-GAPDH (Santa Cruz Biotechnology, #sc-25778, RRID: AB_10167668), anti-tubulin (Santa Cruz Biotechnology, #sc-5286, RRID:AB_628411), anti-BrdU (Abcam, #ab6326, RRID:AB_305426), normal rabbit IgG (Millipore, #12-370, RRID:AB_145841). Secondary antibodies: goat anti-rabbit (Invitrogen, #G-21234, RRID:AB_2536530), goat anti-mouse (Invitrogen, #62-6520, RRID:AB_2533947), donkey anti-rabbit Alexa Fluor 647 (Thermo Fisher, #A-31573, RRID:AB_2536183).

### Reporter gene assay

Reporter gene assays were done as described before (Makkonen et al. 2011). Briefly, COS-1 cells were plated on 12-well plates (120 000 cells per well) and grown for 24 h before changing to stripped medium for 4 h before transfections. COS-1 cells were transfected with 500 ng of p(HRE)_4_tk-TATA-LUC (Tian et al. 2002) along with 100 ng of expression vector encoding for human AR (pSG5-hAR) or human GR (pSG5-hGR). As internal control, the cells were cotransfected with 20 ng of pCMV-beta-galactosidase (ATCC, #77177). As negative control for HRE, the cells were transfected with 500 ng of pGL3-TATA-LUC along with pSG5-hAR or pSG5-hGR as indicated above. After 24 h of transfection, the cells were treated with vehicle (EtOH) or with increasing concentration (100 nM, 1 µM, 10 µM) of Dex, DHT, cortisol (Cort) (Sigma-Aldrich, #H4001), or testosterone (Testo) (Steraloids, #A6950-000) for 18 h. Cell lysis and measurement were performed in triplicates as described (Makkonen et al. 2011). Relative luciferase (LUC) activity was calculated by normalizing with background and beta-galactosidase values. Statistical significance was determined by One-way ANOVA with Bonferroni *post hoc* test.

### Immunoblotting and co-immunoprecipitation (co-IP)

Proteins were isolated and separated as before (Launonen et al. 2021). For hormone treatments, the cells were treated with vehicle (0.1% EtOH), 100 nM Dex, 100 nM DHT or Dex and DHT for 1 h before the start of protein extraction. The following antibodies were utilized: anti-GR (1:1000), anti-AR (1:10000). An anti-GAPDH antibody (1:5000) or anti-tubulin (1:3000) was used as a control for sample loading. Goat anti-rabbit (1:10000) was used as a secondary antibody. Protein bands were detected using Pierce ECL Western Blotting Substrate kit (Thermo Fisher, #32106) and Bio-Rad ChemiDoc Imager. For co-IP, VCaP cells were plated onto 10-cm plates (10 million cells per plate) and cultured in stripped medium for 48 h. The cells were treated with Dex and DHT for 1 h prior to the start of co-IP. The collected cell pellets were resuspended in RIPA-buffer [50 mM Tris-HCl, pH 7.5; 140 mM NaCl; 1 mM EDTA, pH 8.0; 1% Triton X-100; 10% glycerol; 10 mM Na-phosphate, pH 7.0; 10 mM NEM; 1 mM PMSF; 1.5 mM Na_3_VO_4_; 50 mM NaF; 1x protease inhibitor cocktail (PIC, Roche, #11836145001); 1 mM dithiothreitol], sonicated for 10 s, and an aliquot of each supernatant was separated as input sample and combined with equal volume of SDS-PAGE sample buffer complemented with 1xPIC, 10 mM NEM, and 5 % β-mercaptoethanol. Pre-clearing of the remaining supernatants was done with IgG-coupled protein G Dynabeads for 1 h at +4 °C (Invitrogen, #10004D). For immunoprecipitation, supernatants were rotated o/n with either anti-AR (1.5 µl per IP) or IgG (1.5 µl per IP) -coupled magnetic beads at +4 °C. After immunoprecipitation, magnetic beads were washed three times with RIPA-buffer and resuspended into SDS-PAGE sample buffer complemented with 1xPIC, 10 mM NEM, and 5 % β-mercaptoethanol. Samples were heated for 5 min at 95°C and immunoblotted as described above.

### Immunostaining and confocal imaging

VCaP cells were plated on 8-well chamber slides (Ibidi GmbH, #80826) (100 000 cells per well) and cultured in stripped medium for 24 h. Before staining, the cells were treated with vehicle (0.2% EtOH), 100 nM Dex, 100 nM DHT or Dex and DHT for 1 h at 37°C. For fluorescent staining, the cells were fixed with 4% paraformaldehyde in 0.1 M sodium phosphate buffer, pH 7.4 (PB) for 20 min, and washed with PB. The fixed cells were permeabilized for 15 minutes with 0.1% Triton X-100 and 1% BSA, blocked with 1% BSA for 30 min, and incubated o/n at 4°C with anti-AR (1:500) or anti-GR (1:500) antibodies. After washing with PB, the cells were incubated for 2 h with Alexa Fluor 647 labeled secondary antibody (1:500), and nuclei were labeled with 4’,6-diamidino-2-phenylindole (DAPI) (Sigma-Aldrich, #D8417). The fluorescent images were obtained with a Zeiss Axio Observer inverted microscope (40x NA 1.3 oil objective) equipped with a Zeiss LSM 800 confocal module (Carl Zeiss Microscopy GmbH). Image processing was performed using ZEN 3.10 software (Carl Zeiss Microscopy GmbH).

### Cell proliferation assay

For automated live cell proliferation experiments, the cells were seeded onto 96-well plate (30 000 cells per well). On the start of the experiment (day 0), the cells were treated with vehicle (0.2% EtOH), 1 nM Dex, 1 nM DHT or Dex and DHT and the nuclei were stained with IncuCyte NucLight Rapid Red reagent (Essen BioSciences, #4717). The cells were monitored every day for 7 days using IncuCyte S3 Live-Cell Imaging System (Essen BioSciences). IncuCyte S3 2022B software (Essen BioSciences) was used to count the number of cells. Results were normalized to day 0 and are shown as relative percentage in cell count, representing the mean ±SD of 8-10 images per day. Statistical significance was determined by Two-way ANOVA with Bonferroni *post hoc* test.

### ChIP-seq, ATAC-seq and data analysis

Chromatin immunoprecipitation with deep sequencing (ChIP-seq) and assay for transposase-accessible chromatin with sequencing (ATAC-seq) was done as previously described (Helminen et al. 2024). VCaP cells were plated onto 10-cm plates (5 million cells per plate) and grown for 72 h before changing to stripped medium for 48 h. The cells were treated with vehicle (0.1% EtOH), 100 nM Dex, 100 nM DHT or Dex and DHT for 1 h before the start of ChIP or nuclei isolation for ATAC. Antibodies used per IP in ChIP: GR, 2 μg; AR, 3 μl; FOXA1, 2 µg; SMARCA4, 2 µl; BRD4, 2 µl. ChIP-seq sequencing libraries were generated using NEBNext Ultra II DNA Library Prep Kit (New England BioLabs, #E7645L) according to manufacturer’s protocol. Analysis of library quality was done with Agilent 2100 Bioanalyzer using DNA 1000 Analysis kit (Agilent, #5067-1504). In ATAC, isolation of nuclei was verified by Trypan Blue counting. Tn5 transposition reaction was done using 2.5 μl Tagmentase (Diagenode, #C01070012-30). From each sample, 100 000 nuclei were transferred to the reaction. Analysis of ATAC-seq library quality was done with Agilent 2100 Bioanalyzer using High Sensitivity DNA Analysis kit (Agilent, #5067-4626). For ChIP-seq two biological replicate samples were sequenced with Illumina NextSeq 500 (75SE). For ATAC-seq two biological replicate samples were sequenced with Illumina NextSeq 500 (40PE).

ChIP-seq and ATAC-seq read quality filtering and alignment to human hg38 genome was performed as previously described (Helminen et al. 2024). Downstream data analysis was performed using HOMER (RRID:SCR_010881) (Heinz et al. 2010) as previously described (Helminen et al. 2024). For GR-binding sites, differential peak clusters were determined using getDifferentialPeaksReplicates.pl between the single and dual hormone treatments. In C1, dual hormone treated sample (DexDHT) differed from Dex treated sample (FDR < 0.05, fold change > 2). In C2, dual hormone treated sample did not differ from Dex treated sample. In C3, Dex treated sample differed from DexDHT treated sample (FDR < 0.05, fold change < 2). C3 represents <100 binding sites and was left out of the downstream analyses. Division of AR-binding sites based on preaccessibility to quartiles (Q1-4) was performed with annotatePeaks.pl using ATAC-seq signal from EtOH treated sample. Q1 sites were the most and Q4 the least preaccessible. All peaks are presented in Supplemental Table S1. Heatmaps were generated with 20 bp bins surrounding ±1 kb area around the center of the peak. All plots were normalized to 10 million mapped reads and further to local tag density, tags per bp per site. Box plots and scatter plots represented log_2_ tag counts. De novo motif searches were performed using findMotifsGenome.pl with the following parameters: 200 bp peak size window, strings with 2 mismatches, binomial distribution to score motif *p*-values, and 50 000 background regions. Full motif analysis data is presented in Supplemental Table S2. For motif scatter plot, only motifs with over 10% target sequence enrichment and *p* < 0.01 in at least one condition were included. Scatter plot displays log_2_ fold change of motif enrichment *p*-values between C1 and C2 sites. Motifs with log_2_ fold change ±0.5 were considered C1 or C2 preferred motifs. Motif score and HRE count analyses were performed with annotatePeaks.pl using the following HOMER motifs: FOXA1 (LNCAP); HOXB13 (ProstateTumor); Gata2(K562); Nkx3.1 (LNCaP); ARE (LNCAP); GRE (A549); PR (T47D); AR-halfsite (LNCaP). Box plots were drawn with Tukey method with statistical significance determined by unpaired *t*-test (two conditions) or One-way ANOVA with Bonferroni *post hoc* test (three or more conditions).

### ChIP- and sequential (re)ChIP-qPCR

ChIP was performed as indicated above. Briefly, the cells were plated onto 10-cm plates (22Rv1, 3 million cells per plate; PC3-AR, 1.5 million cells per plate) and grown for 72 h before changing to stripped medium for 48 h. The cells were treated with vehicle (0.1% EtOH), 100 nM Dex, 100 nM DHT or Dex and DHT for 1 h before the start of ChIP. Antibodies used per IP in ChIP: GR, 2 µg (for #sc-1003) or 12.5 μl (for #12041); AR, 3 μl. For reChIP performed in VCaP cells, after the first ChIP the immunocomplexes were eluded from magnetic beads using 10 mM dithiothreitol. Subsequently, the second ChIP was performed as indicated above. Antibodies used per IP in the first ChIP: GR, 2 µg. Antibodies used per IP in the second ChIP: AR, 3 µl; FOXA1, 2 µg. The qPCR was performed using LightCycler 480 SYBR Green I Master (Roche, #04887352001) and LightCycler 480 Instrument II (Roche) and the binding was calculated with formula E^-(ΔCt)^ x 10, where E (efficiency of target amplification) is a coefficient of DNA amplification by one PCR cycle for a particular primer pair and ΔCt is Ct_(ChIP template)_ - Ct_(Input)_. Primers used are presented in Supplemental Table S5. AR and GR do not bind to *KLK3* site in PC3-AR cells.

### GRO-seq and data analysis

Global run-on sequencing (GRO-seq) libraries were done as previously described (Toropainen et al. 2016). VCaP cells were plated onto 15-cm plates (12.5 million cells per plate) and grown for 72 h before changing to stripped medium for 48 h. The cells were treated with vehicle (0.1% EtOH), 100 nM Dex, 100 nM DHT or Dex and DHT for 1 h before the start of nuclei isolation for GRO-seq. After isolation, 5 million nuclei were used for run-on reactions in the presence of Br-UTP (Santa Cruz Biotechnology, #sc-214314A), and RNA was isolated using TRIZON-LS reagent (Invitrogen, #10296028). Labeled RNA was purified with anti-BrdU conjugated to Protein G Dynabeads (Invitrogen, #10004D). After polyA-tailing, cDNA synthesis and circularization, samples were PCR amplified and size-selected on 10% TBE gels. Two biological replicate samples were sequenced with Illumina HiSeq 2000 (50SE).

GRO-seq reads were quality controlled and trimmer with FASTX-toolkit (RRID:SCR_005534) and HOMER. Reads were aligned to human hg38 genome using bowtie (RRID:SCR_005476) (Langmead et al. 2009) reporting only reads with maximum 2 mismatches and keeping only the best aligned read. Differentially expressed genes were analyzed with DESeq2 (RRID:SCR_015687) (Love et al. 2014) through HOMER (Heinz et al. 2010) for all comparisons. Genes differentially expressed between hormone treatments were defined with criteria FDR <0.01 and log_2_ fold change ±0.5 between differently treated samples. All GRO-seq data values are presented in Supplemental Table S3. For analysis of enhancer RNA (eRNA) only intergenic binding sites were used. Histograms and box plots were generated with annotatePeaks.pl limiting the number of reads at each unique position to three. Histograms display enrichment of eRNA in strand-specific manner, and box plots were generated using the Tukey method. Definition of synergistically DexDHT-induced genes were done as previously described (Goldberg et al. 2022). Briefly, for each DHT and DexDHT-induced gene EtOH fragments per kilobase per million (FPKM) value was reduced from the single or dual hormone treatment FPKM value. Subsequently, reduced single hormone treatment values were added together. Finally, if the reduced DexDHT value divided by the added single hormone values was over 1, the gene was designated as synergistically DexDHT-induced gene. If the value was less than 1, the gene was designated as DHT- and DexDHT-induced gene. For simplicity, DHT- and DexDHT-induced genes are indicated as non-synergistic, and synergistically DexDHT-induced genes are indicated as synergistic. List of non-synergistic and synergistic genes are presented in Supplemental Table S4. Gene-to-peak association (gene-centric analysis) was performed using annotatePeaks.pl measuring the linear distance from target gene TSS to center of the peak. Here, C1 peaks were linked to the nearest synergistic or non-synergistic genes, independent of C2 peaks, while C2 peaks were similarly associated, regardless of C1 peaks. Statistical significance was determined with One-way ANOVA with Bonferroni *post hoc* test. For pathway analyses, the different gene clusters were enriched to Hallmark Gene Sets using Metascape (RRID:SCR_016620) (Zhou et al. 2019), with criteria *p* < 0.01, at least three gene overlap per gene set, and >1.5 pathway enrichment. Enriched pathways are presented in Supplemental Table S4.

### RNA extraction and RT-qPCR

For RNA isolation, the cells were plated onto 12-well plates (VCaP, 350 000 cells per well; PC3-AR, 100 000 cells per well) and grown for 72 h before changing to stripped medium for 48 h. The cells were treated with vehicle (0.1% EtOH), 100 nM Dex, 100 nM DHT or Dex and DHT for 18 h before the start of RNA extraction. Total RNA was extracted using TriPure Isolation Reagent (Roche, #1166715700). Quality and quantity of the isolated RNA were determined with NanoDrop One (Thermo Scientific). 1 μg of total RNA from each sample was converted into cDNA with First Strand cDNA Synthesis Kit (Roche, #04897030001). The qPCR was performed using LightCycler 480 SYBR Green I Master (Roche, #04887352001) and LightCycler 480 Instrument II (Roche) and the relative gene expression was calculated using the 2^−(ΔΔCt)^ method, where ΔΔCt is ΔCt_(treatment)_ - ΔCt_(vehicle)_ and ΔCt is Ct_(target gene)_ - Ct_(control gene)_. *RPL13A* was used as a control gene. Primers used are presented in Supplemental Table S5.

### scRNA-seq and data analysis

For single-cell RNA sequencing (scRNA-seq), VCaP cells were plated onto 10-cm plates (10 million cells per plate) and grown for 72 h before changing to stripped medium for 48 h. The cells were treated with vehicle (0.1% EtOH), 100 nM Dex, 100 nM DHT or Dex and DHT for 18 h before the start of extraction. Single cell suspensions were harvested and washed according to published protocol (10x Genomics, #CG00054). Isolated cells were concentrated to 700 cells per µl. scRNA-seq libraries were generated using Single Cell 3’ Kit v3.1 (10x Genomics, #PN-1000269, #PN-1000127, #PN-1000215) according to published protocol (10x Genomics, #CG000315). Target cell recovery was set at 2000 cells. Analysis of scRNA-seq library quality was done with Agilent 2100 Bioanalyzer using High Sensitivity DNA Analysis kit (Agilent, #5067-4626). scRNA-seq libraries were sequenced with Illumina NextSeq 2000 (100PE).

ScRNA-seq reads were mapped to a reference genome hg38 (GRCh38-2020-A) using Cell Ranger 7.0.0 (RRID:SCR_017344). In total 12352 cells were obtained from sequencing the EtOH (2093 cells), Dex (2410 cells), DHT (4052 cells), and DexDHT (3797 cells) treated VCaP cell fractions. DecontX was used to estimate and remove ambient RNA related contamination (Yang et al. 2020). Filtering of cells was based on presence of duplicates and deficient counts of features. Deficient cells had more than 4000 (Dex and DHT) or 6000 (EtOH and DexDHT) or less than 750 features. Additionally, genes expressed in less than three individual cells were removed, and cells with high mitochondrial reads (over 40%) and low ribosomal reads (less than 3%) were filtered out. After filtering, 4956 cells were retained from EtOH (1392), Dex (1110), DHT (1194), and DexDHT (1260) treated VCaP cells. Further downstream analysis was conducted in R 4.2.2 (RRID:SCR_001905) using Seurat 4.3.0 (RRID:SCR_007322) (Hao et al. 2021). The pre-processed matrices from each sample were merged, and SCTransform function was applied performing normalization, scaling, feature selection, and regressing out variation due to mitochondrial expression (Hafemeister and Satija 2019). UMAP dimensional reduction was generated with RunUMAP function using 30 principal components. The nearest neighbor graph was generated with default settings using FindNeighbors command with 30 principal components. Cell subpopulations were identified with FindClusters function using default Louvain algorithm. Visualization of data was performed with Seurat. Data in UMAP is represented as mRNA expression (log1p normalized). Data in dot and violin plots is represented as the average of the normalized expression values. Here, scaled expression values center the expression to have zero mean and SD of one. Gene-to-peak association (gene-centric analysis) was performed using annotatePeaks.pl with C1 GRBs and differentially expressed genes between DHT and DexDHT scRNA-seq samples (*p* < 0.01 and log_2_ fold change >0.25) regardless of subpopulations. The number of cells expressing a specific gene was performed from the filtered cells. The specific gene in a particular cell had to have non-zero value to be considered expressed. Pathway enrichment was done as described for GRO-seq data, and enriched pathways are presented in Supplemental Table S4.

### Public datasets and patient data analysis

The following publicly available sequencing datasets were used: GSE136016 for AR and SMARCA4 ChIP-seq in EtOH and DHT exposed VCaP cells (Launonen et al. 2021). GSE214757 for FOXA1 ChIP-seq in EtOH, Dex and DHT exposed VCaP cells, and AR ChIP-seq and ATAC-seq in enzalutamide (ENZ) exposed VCaP cells (Helminen et al. 2024). GSE256370 for RNA-seq in EtOH, estradiol (E2), R1881 (synthetic androgen) and E2+R1881 exposed VCaP cells (Lafront et al. 2024). The data was processed as indicated above and as previously described (Helminen et al. 2024). PCa patient gene expression and survival data were obtained through cBioPortal (RRID:SCR_014555) (Cerami et al. 2012), Xena (RRID:SCR_018938) (Goldman et al. 2020), GEPIA2 (RRID:SCR_018294) (Tang et al. 2019), and Prostate Cancer Atlas (Bolis et al. 2021). For patient survival and box plots, the patients were divided based on median to low and high expressing populations. Statistical significance was calculated with log-rank test for survival, and with One-way ANOVA with Bonferroni *post hoc* test for box plots.

## DATA ACCESS

All raw and processed sequencing data generated in this study have been submitted to the NCBI Gene Expression Omnibus (GEO; https://www.ncbi.nlm.nih.gov/geo/) under accession number GSE266217.

## ACKNOWLEDGEMENTS

We thank Merja Räsänen and Saara Pirinen for technical assistance. The EMBL GeneCore team (Heidelberg, Germany) is greatly acknowledged for deep sequencing services. This work was carried out with the support of UEF Single Cell Genomics Core and UEF Cell and Tissue Imaging Unit, University of Eastern Finland, Finland, Biocenter Kuopio and Biocenter Finland. The computational analyses were performed on servers provided by UEF Bioinformatics Center, University of Eastern Finland, Finland. This work was supported by the Research Council of Finland; Cancer Foundation Finland; Sigrid Jusélius Foundation; University of Eastern Finland strategic funding; University of Eastern Finland Doctoral Programme of Molecular Medicine.

## AUTHOR CONTRIBUTIONS

**Johannes Hiltunen**: Formal Analysis; Investigation; Visualization; Writing – review & editing. **Laura Helminen**: Formal Analysis; Investigation; Visualization; Writing – review & editing. **Niina Aaltonen**: Formal Analysis; Investigation; Visualization; Writing – review & editing. **Kaisa-Mari Launonen**: Investigation; Writing – review & editing. **Hanna Laakso**: Investigation; Writing – review & editing. **Marjo Malinen**: Investigation; Writing – review & editing. **Einari A. Niskanen**: Formal Analysis; Investigation; Writing – review & editing. **Jorma J. Palvimo**: Funding acquisition; Supervision; Writing – review & editing. **Ville Paakinaho**: Conceptualization; Formal Analysis; Funding acquisition; Investigation; Project administration; Supervision; Visualization; Writing – original draft; Writing – review & editing.

