## Supplemental material for "Androgen receptor-mediated assisted loading of the glucocorticoid receptor modulates transcriptional responses in prostate cancer cells"

### Table of Contents

- **Supplemental Fig. S1**
- **Supplemental Fig. S2**
- **Supplemental Fig. S3**
- **Supplemental Fig. S4**
- **Supplemental Fig. S5**
- **Supplemental Fig. S6**
- **Supplemental Fig. S7**
- **Supplemental Table legends**

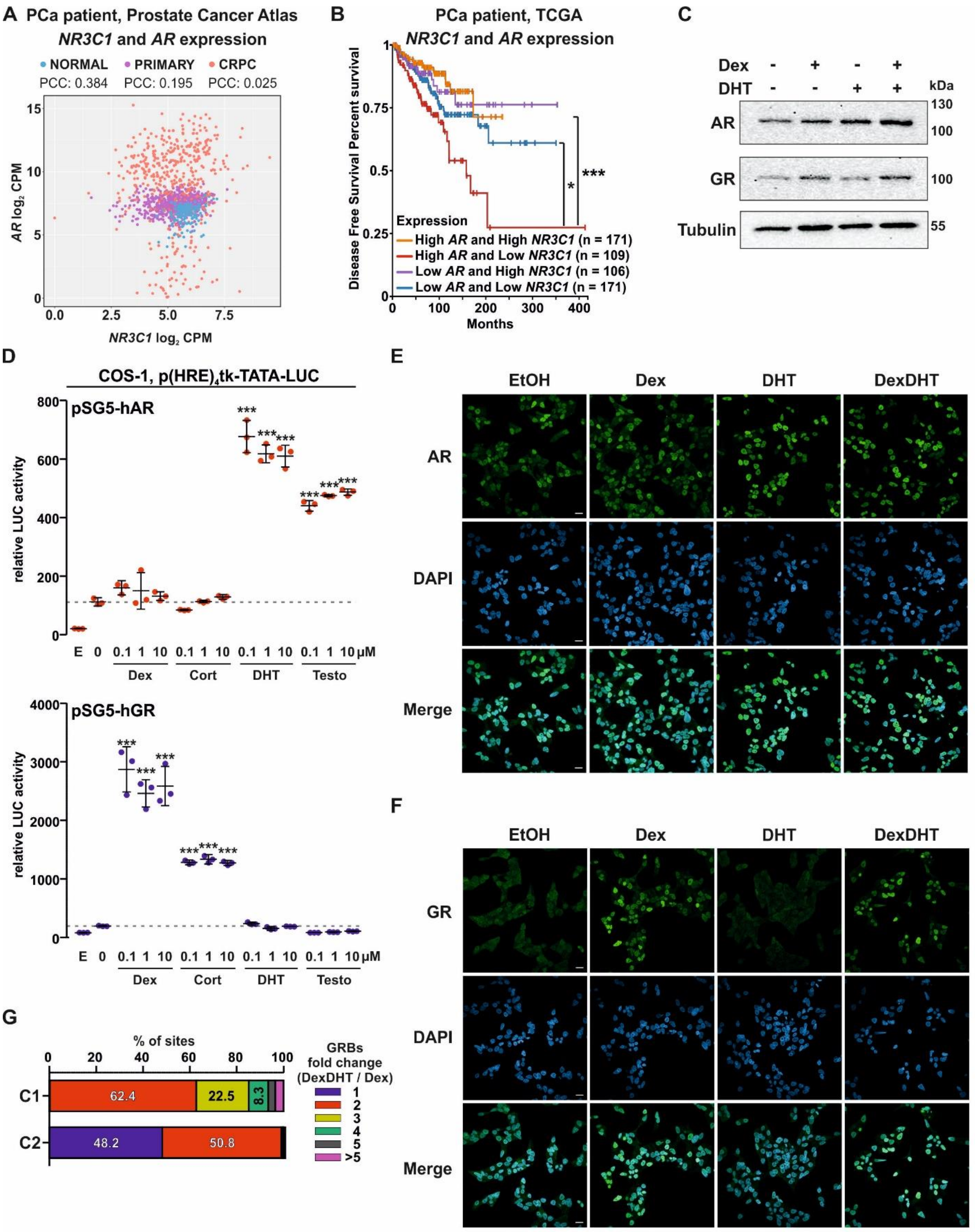


**Supplemental Fig. S1. Relationship of *AR* and *NR3C1* expression in prostate cancer patients.** (**A**) Scatter plot of *NR3C1* (x-axis) and *AR* (y-axis) expression from RNA-seq data from Prostate Cancer Atlas patient datasets. Data represent normal, primary cancer, and castration resistant cancer (CRPC). Correlation displayed as Pearson Correlation Coefficiency (PCC). (**B**) Disease-free survival of patients with high or low expression of *AR* and *NR3C1* in TCGA dataset: high *AR* and high *NR3C1* (orange); high *AR* and low *NR3C1* (red); low *AR* and high *NR3C1* (purple); low *AR* and low *NR3C1* (blue). Statistical significance calculated with log-rank test. (**C**) Immunoblotting of AR, GR and tubulin protein levels in EtOH, Dex, DHT, and DexDHT treated VCaP cells. (**D**) Reporter gene assay from COS-1 cells transfected with p(HRE)$_4$tk-TATA-LUC with pSG5-hAR (upper) or pSG5-hGR (lower), and treated with indicated µM concentration of Dex, Cort, DHT, and Testo. Control cells (marked as E) were transfected with pGL3-TATA-LUC. Line and whiskers represent average ± SD, $n$=3. Statistical significance calculated using One-way ANOVA with Bonferroni *post hoc* test. (**E-F**) Immunofluorescence (IF) images of (**E**) AR and (**F**) GR from VCaP cells. The cells were treated with EtOH, Dex, DHT, and DexDHT. DAPI was used as a marker for nuclei. Scale bar, 20 µm. (**G**) Percentage of C1 and C2 sites displaying specific fold change (FC) difference in GR ChIP-seq between DexDHT and Dex treatments: FC no change (1), blue; FC at least 2, red; FC at least 3, yellow; FC at least 4, green; FC at least 5, dark grey; FC over 5, magenta. *, $p < 0.05$; ***, $p < 0.001$.

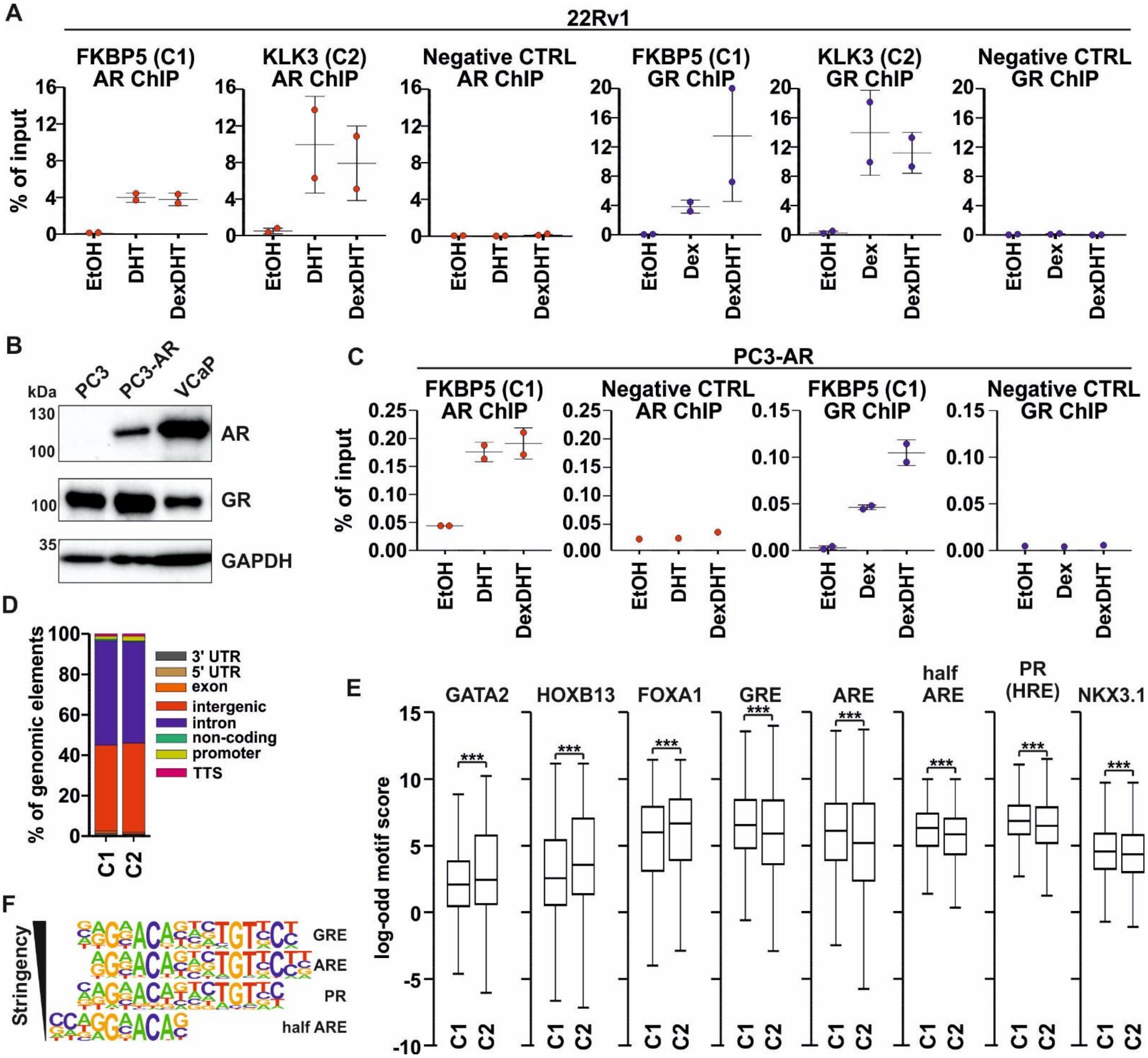


**Supplemental Fig. S2. AR-mediated assisted loading of GR in 22Rv1 and PC3-AR cells.** (**A**) ChIP-qPCR analysis of AR (upper) and GR (lower) binding at *FKBP5*, *KLK3*, or negative CTRL loci in EtOH, Dex (GR only), DHT (AR only), and DexDHT treated 22Rv1 cells. (**B**) Immunoblotting of AR, GR and GAPDH protein levels in PC3, PC3-AR, and VCaP cells. (**C**) ChIP-qPCR analysis of AR (left) and GR (right) binding at *FKBP5* or negative CTRL loci in EtOH, Dex (GR only), DHT (AR only), and DexDHT treated PC3-AR cells. AR and GR do not bind to *KLK3* loci in PC3-AR cells. Line and whiskers represent average ± SD, *n*=1-2. (**D**) Percentage of C1 and C2 peak enrichment at genomic elements: 3'UTR (grey); 5'UTR (light brown); exon (orange); intergenic (red); intron (blue); non-coding (green); promoter (yellow); TTS (magenta). (**E**) Motif score of GATA2, HOXB13, FOXA1, GRE, ARE, half ARE, PR, and NKX3.1 motif enrichment at C1 and C2 peaks. Statistical significance calculated with unpaired *t*-test. (**F**) Position weight matrix (PWM) logos for GRE, ARE, PR, and half ARE motifs used by HOMER program. The motif stringency based on consensus bases are indicated with black triangle.

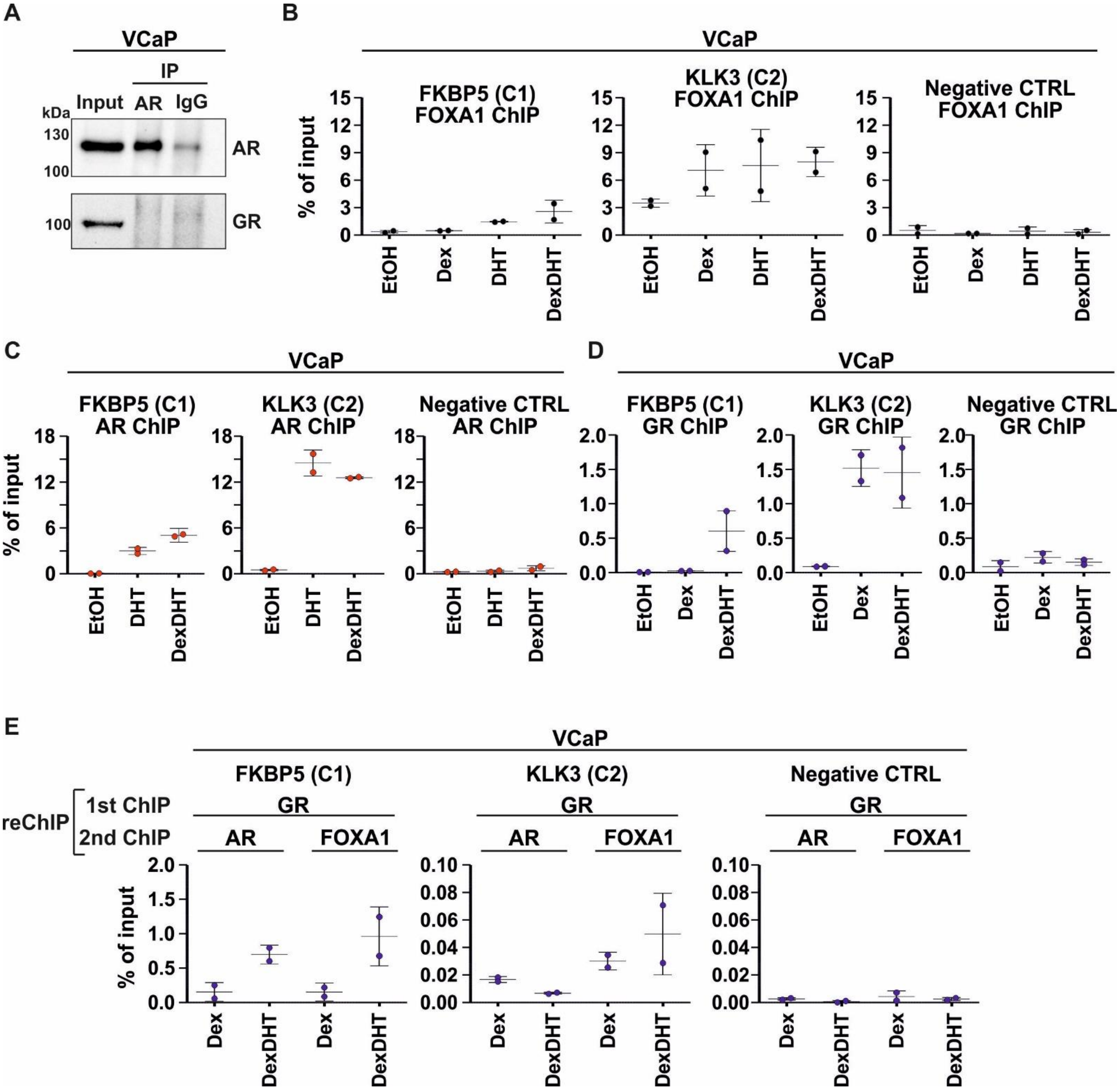


**Supplemental Fig. S3. Sequential GR chromatin binding with AR and FOXA1 in VCaP cells.** (**A**) Whole-cell lysates (Input), AR and normal rabbit IgG immunoprecipitated (IP) samples were immunoblotted with AR or GR antibodies. (**B**) ChIP-qPCR analysis of FOXA1 binding at *FKBP5*, *KLK3*, or negative CTRL loci in EtOH, Dex, DHT, and DexDHT treated VCaP cells. (**C**) ChIP-qPCR analysis of AR binding at *FKBP5*, *KLK3*, or negative CTRL loci in EtOH, DHT, and DexDHT treated VCaP cells. (**D**) ChIP-qPCR analysis of GR binding at *FKBP5*, *KLK3*, or negative CTRL loci in EtOH, Dex, and DexDHT treated VCaP cells. (**E**) Sequential (re)ChIP-qPCR analysis of GR+AR or GR+FOXA1 binding at *FKBP5*, *KLK3*, or negative CTRL loci in Dex and DexDHT treated VCaP cells. Line and whiskers represent average ± SD, *n*=2.

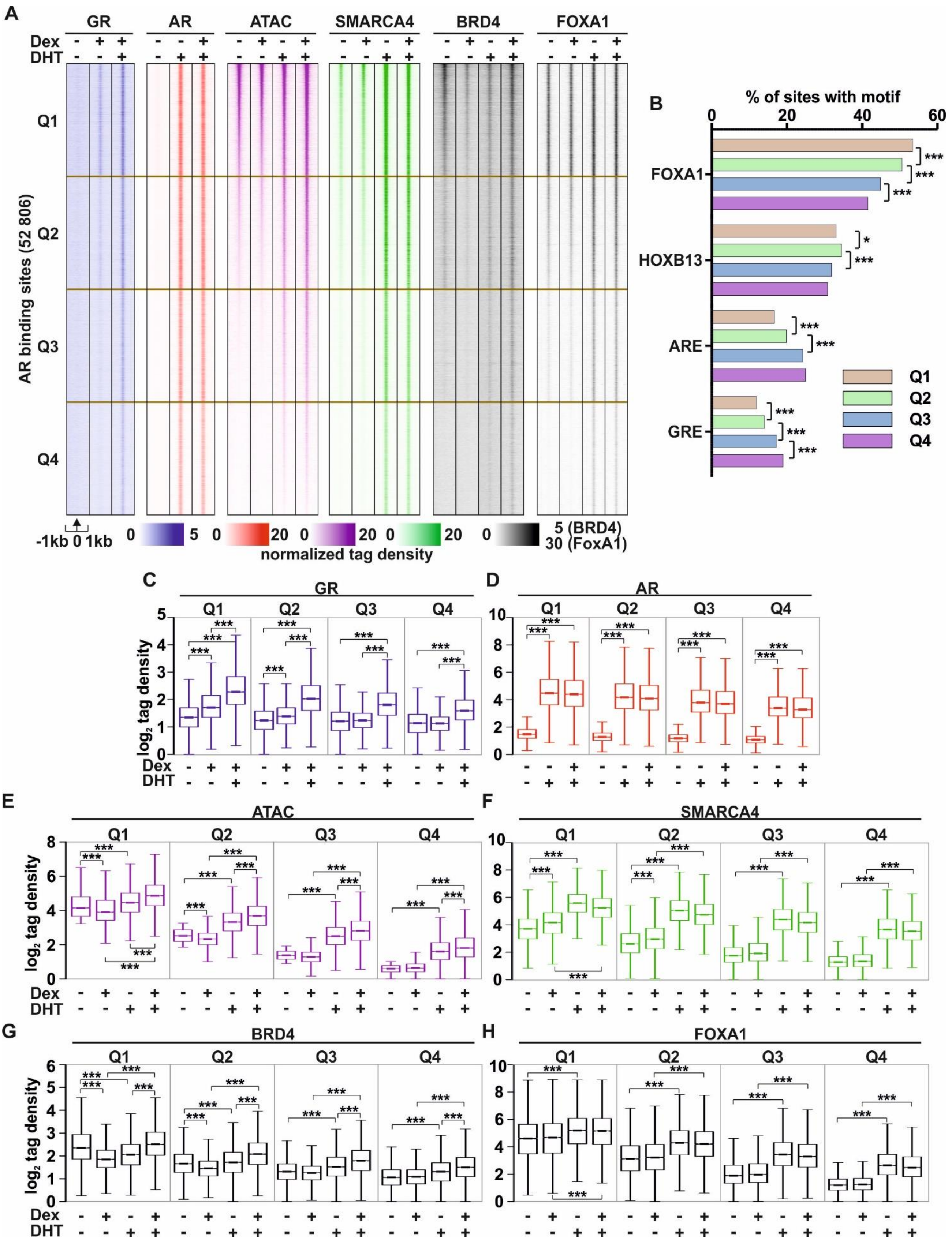


**Supplemental Fig. S4. Analysis of GR and AR binding to site with variable preaccessibility.** (**A**) GR ChIP-seq, AR ChIP-seq, ATAC-seq, SMARCA4 ChIP-seq, BRD4 ChIP-seq, and FOXA1 ChIP-seq profiles at Q1, Q2, Q3, and Q4 sites in VCaP cells. Q1-Q4 represent ARBs divided into quartiles based on chromatin preaccessibility. Q1 are the most and Q4 the least pre-accessible sites. Each heatmap represents ± 1 kb around the center of the GR peak. Binding intensity (tags per bp per site) scale is noted below on a linear scale. (**B**) Percentage of FOXA1, HOXB13, ARE, and GRE motif enrichment at Q1 (brown bars), Q2 (green bars), Q3 (blue bars) and Q4 (purple bars) peaks. Statistical significance calculated with Chi-squared test. (**C-H**) Box plots represent the normalized $log_2$ tag density of (**C**) GR ChIP-seq, (**D**) AR ChIP-seq, (**E**) ATAC-seq, (**F**) SMARCA4 ChIP-seq, (**G**) BRD4 ChIP-seq, (**H**) FOXA1 ChIP-seq at the Q1-Q4 sites. Statistical significance calculated using One-way ANOVA with Bonferroni *post hoc* test. All heatmaps and box plots are normalized to a total of 10 million reads. *, $p < 0.05$; ***, $p < 0.001$.

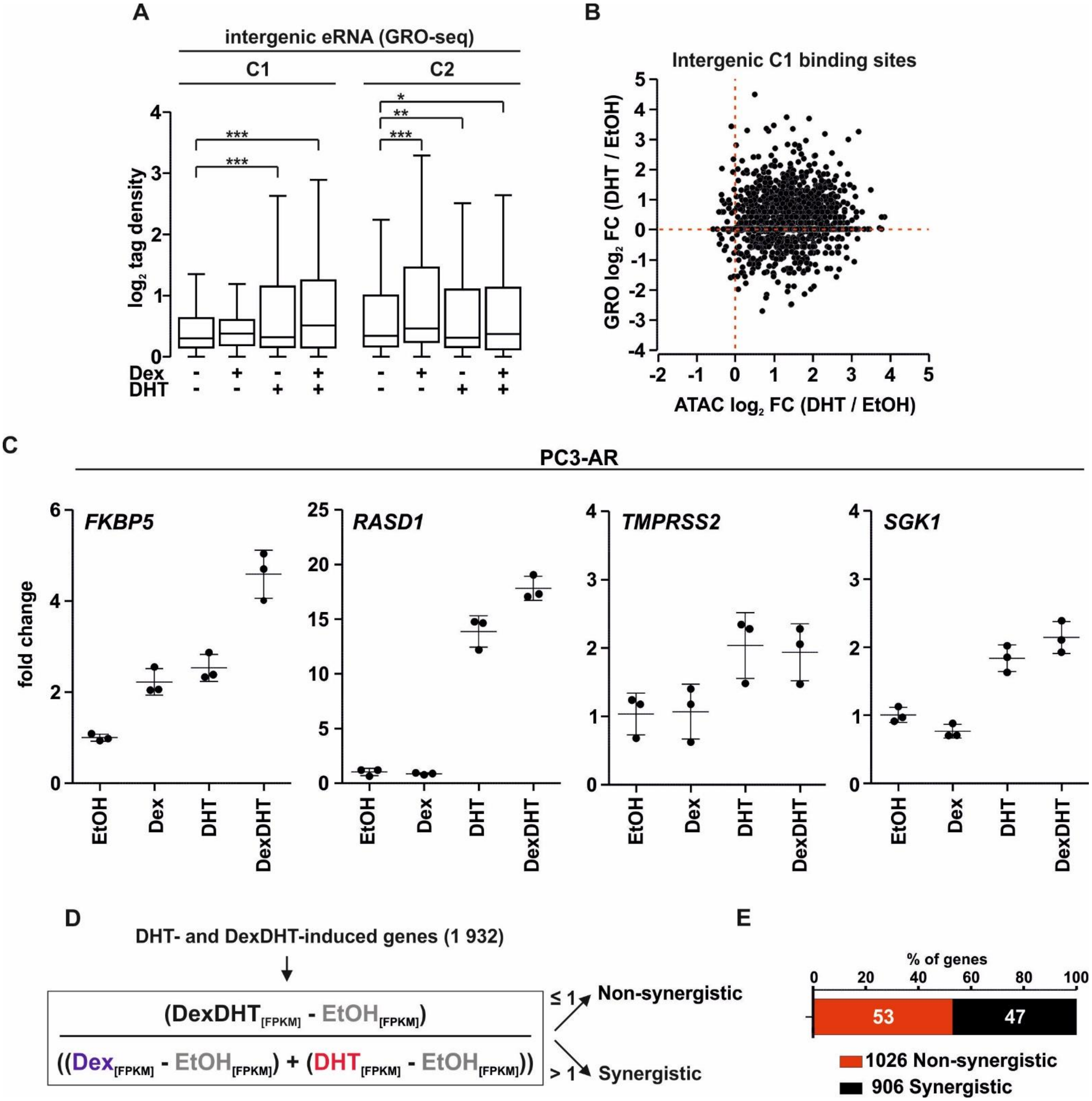


**Supplemental Fig. S5. Definition of synergistic DexDHT-induced genes.** (**A**) Box plots represent the normalized $\log_2$ tag density of GRO-seq tags at intergenic C1 and C2 peaks. Statistical significance calculated using One-way ANOVA with Bonferroni *post hoc* test. Box plots are normalized to a total of 10 million reads. (**B**) Scatter plot of ATAC-seq (x-axis) and GRO-seq (y-axis) $\log_2$ fold change (FC) between DHT and EtOH treated samples at intergenic C1 sites. (**C**) RT-qPCR analysis of *FKBP5*, *RASD1*, *TMPRSS2*, or *SGK1* expression in EtOH, Dex, DHT, and DexDHT treated PC3-AR cells. Line and whiskers represent average ± SD, $n$=3. (**D**) Formula used to define DHT- and DexDHT-induced and synergistically DexDHT-induced genes. (**E**) Bar graph depicts the number of DHT- and DexDHT-induced genes that are defined as DHT- and DexDHT-induced (red color) and synergistically DexDHT-induced (black color) genes. All box plots are normalized to a total of 10 million reads. *, $p < 0.05$; **, $p < 0.01$; ***, $p < 0.001$.

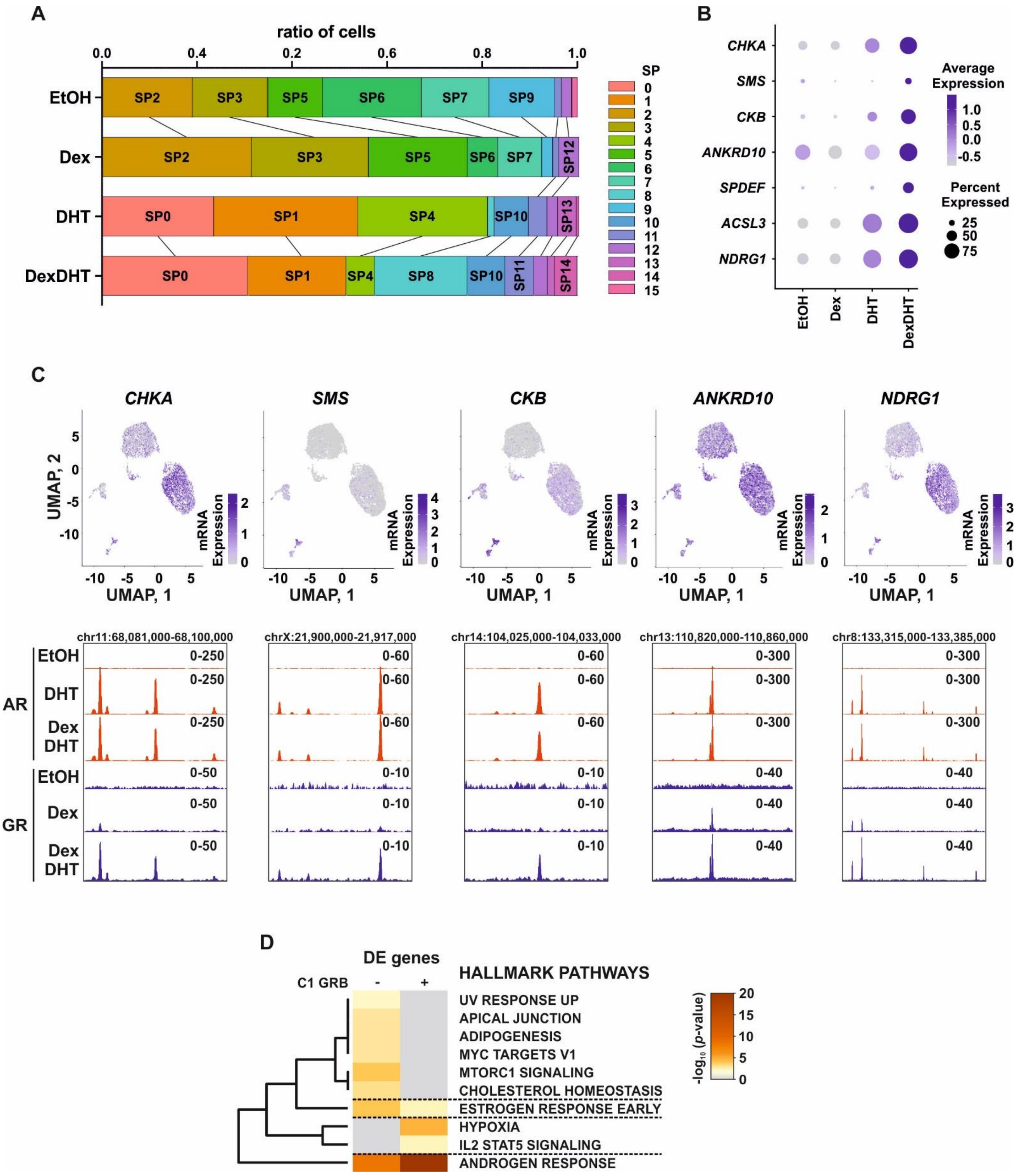


**Supplemental Fig. S6. Ratio of cells at single cell clusters changes upon treatments.** (**A**) Bar graph depicts the ratio of single cells at subpopulations 0-14 (SP0-SP14) defined with UMAP from scRNA-seq. Clusters are depicted with distinct color and label. (**B**) Dot plot of the expression of selected genes in single cells based on treatments. Intensity of the color depicts the average expression, and size of the circle depicts the percentage of cells that express the gene. (**C**) UMAP of gene expression at the single cell clusters (upper) and genome browser tracks (lower) of chromatin binding of AR and GR for *CHKA*, *SMS*, *CKB*, *ANKRD10*, and *NDRG1*. The intensity of the color depicts the mRNA expression (log1p normalized). (**D**) Hallmark Gene Sets pathway analysis of differentially expressed (DE) genes between DHT and DexDHT in scRNA-seq data with (+) or without (-) association with C1 GRB. Color scale represents -$\log_{10}$ *p*-value. All genome browser tracks are normalized to a total of 10 million reads.

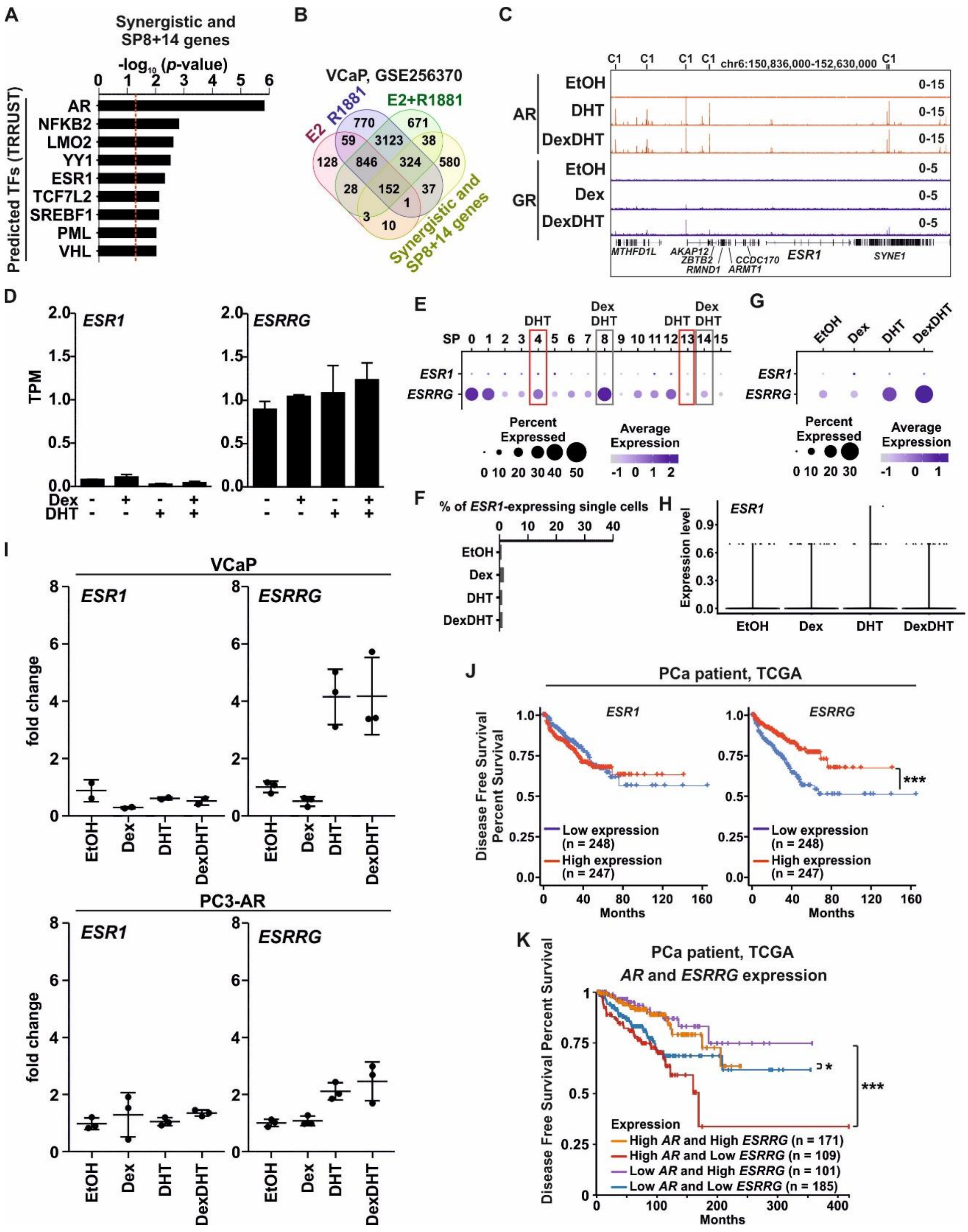


**Supplemental Fig. S7. *ESR1* and *ESRRG* expression in prostate cancer.** (**A**) TRRUST regulatory TFs prediction analysis from synergistic and subpopulation (SP)8+14 genes. Bar graphs depict the -$\log_{10}$ *p*-value of the enrichment with red dashed line indicating p=0.05. (**B**) Venn diagrams depict the overlap of estradiol (E2) (red circle), R1881 (blue circle), and E2+R1881 (green circle) regulated genes in VCaP cells with synergistic and SP8+14 genes (yellow circle). (**C**) Genome browser track depicts chromatin binding of AR and GR at the *ESR1* locus. (**D**) Bar graphs depict the expression of *ESR1* (left) and *ESRRG* (right) using GRO-seq tags (TPM values). (**E**) Dot plot depicts the expression *ESR1* and *ESRRG* at single cell subpopulations 0-15 (SP0-SP15) in VCaP cells. Intensity of the color depicts the average expression, and size of the circle depicts the percentage of cells that express the gene. DHT dominated SPs are highlighted with red rectangle, and DexDHT dominated SPs are highlighted with grey rectangle. (**F**) Percentage of *ESR1*-expressing single cells in EtOH, Dex, DHT, and DexDHT treated VCaP cells. (**G**) Dot plot of the expression of *ESR1* and *ESRRG* in single cells based on treatments. Intensity of the color depicts the average expression, and size of the circle depicts the percentage of cells that express the gene. (**H**) Violin plot represents the single cell expression level of *ESR1* in EtOH, Dex, DHT, and DexDHT treated VCaP cells. (**I**) RT-qPCR analysis of *ESR1* or *ESRRG* expression in EtOH, Dex, DHT, and DexDHT treated VCaP (upper) and PC3-AR (lower) cells. Line and whiskers represent average ± SD, $n$=2-3. (**J**) Disease-free survival of TCGA PCa patients with high (red) or low (blue) expression levels of *ESR1* (left) and *ESRRG* (right). Statistical significance calculated with log-rank test. (**K**) Disease-free survival of TCGA PCa patients with combination of high and/or low expression levels of *AR* and *ESRRG*. Statistical significance calculated with log-rank test. All genome browser tracks are normalized to a total of 10 million reads. *, $p < 0.05$; ***, $p < 0.001$.

### Supplemental Table legends

**Supplemental Table S1. ChIP-seq peak clusters.** GR-binding sites, C1-C3, and AR-binding sites, Q1-Q4.

**Supplemental Table S2. Motif analysis data from peak clusters.** GR-binding sites, C1-C2, and AR-binding sites, Q1-Q4.

**Supplemental Table S3. GRO-seq analyses.** DESeq2 results and TPM values.

**Supplemental Table S4. Gene lists and pathway analyses.** List of synergistic and non-synergistic genes, and differentially DexDHT-regulated genes in scRNA-seq. Pathway enrichments from Metascape.

**Supplemental Table S5. List of qPCR primers.** Primers used in RT-qPCR and ChIP-qPCR.
